# DdiA, an XRE family transcriptional regulator, regulates a LexA-independent DNA damage response in *Myxococcus xanthus*

**DOI:** 10.1101/2025.02.19.639066

**Authors:** Jana Jung, Timo Glatter, Marco Herfurth, Lotte Søgaard-Andersen

## Abstract

Repair of DNA damage is essential for genome integrity. DNA damage elicits a DNA damage response (DDR) that includes error-free and error-prone, i.e. mutagenic, repair. The SOS response is a widely conserved system in bacteria that regulates the DDR and depends on the recombinase RecA and the transcriptional repressor LexA. However, RecA/LexA-independent DDRs have been identified in several bacterial species. Here, using a whole-cell, label-free quantitative proteomics approach, we map the proteomic response in *Myxococcus xanthus* to mitomycin C treatment and the lack of LexA. In doing so, we confirm a LexA-independent DDR in *M. xanthus*. Using a candidate approach, we identify DdiA, a transcriptional regulator of the XRE family, and demonstrate that it regulates a subset of the LexA-independent DDR genes. *ddiA* is expressed heterogeneously in a subpopulation of cells in the absence of exogenous genotoxic stress and reversibly induced population-wide in response to such stress. DdiA, indirectly or directly, activates the expression of *dnaE2*, which encodes the DnaE2 error-prone DNA polymerase, and inhibits the expression of *recX*, which encodes RecX, a negative regulator of RecA. Accordingly, the Δ*ddiA* mutant has a lower mutation frequency than the wild-type but also a fitness defect, suggesting that DdiA mediates a trade-off between fitness and mutagenesis. We speculate that the DdiA-dependent response is tailored to counter replication stress, thereby preventing the induction of the complete RecA/LexA-dependent DDR in the absence of exogenous genotoxic stress.

**Importance:** DNA damage repair is essential for genome integrity and depends on the DNA damage response (DDR). While the RecA/LexA-dependent SOS response is widely conserved in bacteria, there are also RecA/LexA-independent DDRs. Here, we identify the DNA damage-induced transcriptional regulator DdiA in *Myxococcus xanthus* and demonstrate that it regulates part of the RecA/LexA-independent DDR. DdiA activates the expression of *dnaE2*, which encodes the DnaE2 error-prone DNA polymerase, and inhibits the expression of *recX*, which encodes RecX, a negative regulator of RecA. Because the Δ*ddiA* mutant has a lower mutation frequency than the wild-type but also a fitness defect, we suggest that DdiA mediates a trade-off between fitness and mutagenesis and that the DdiA-dependent DDR is specifically tailored to counter replication stress.

## Introduction

In their natural environments, bacteria are exposed to exogenous genotoxic stress, including radiation, UV light, toxins, antibiotics, and other chemicals (1–5). DNA damage also occurs spontaneously due to endogenous factors such as reactive oxygen and nitrogen species, which are byproducts of metabolism, and replication stress (1, 2, 4, 5). Irrespective of the source, DNA damage is a threat to genome integrity and cellular survival; however, DNA damage also helps generate the genetic variation that is key to evolutionary changes (6).

DNA damage elicits a two-pronged DNA damage response (DDR) in bacteria. One part involves the synthesis of conserved enzymes that execute homologous recombination and error-free DNA repair to remedy the DNA damage (2, 3). The second part involves the synthesis of low-fidelity DNA polymerases that carry out error-prone, i.e. mutagenic, translesion synthesis (TLS), thereby enabling replication fork progression past damaged DNA (4, 7). While the first part helps to repair the damage and maintain genome integrity, the second part significantly contributes to DNA damage-induced mutagenesis (6). The DDR can also activate cell cycle checkpoints that delay cell division until the damage has been repaired (3, 8).

The SOS response is a widely conserved system in bacteria that controls the DDR and relies on two key regulators: the recombinase RecA and the transcriptional regulator LexA (2, 3). In the absence of DNA damage, LexA binds to the promoter regions of its target genes to repress transcription. In response to DNA damage, single-stranded DNA (ssDNA) accumulates and binds to RecA. The RecA/ssDNA complex interacts with LexA and acts as a co-protease, stimulating autocleavage of LexA, resulting in the derepression of genes for DNA damage repair and error-prone DNA repair. The *lexA* gene is part of the LexA regulon, and LexA is a negative autoregulator; the increased LexA synthesis during the SOS response helps to ensure the repression of the SOS genes once the repair processes are completed. However, LexA is not universally conserved in bacteria (2, 3, 9). Likewise, the specific DNA repair genes regulated by the RecA/LexA system vary between species (1–3). Accordingly, RecA/LexA-independent DDRs have been identified in several bacterial species (10–15).

*Mycobacterium smegmatis* and *M. tuberculosis* have a RecA/LexA-dependent DDR, but the heterodimeric transcriptional activator PafBC is the key regulator of the DDR and directly activates the expression of numerous DNA repair genes independently of RecA/LexA (16–19). More recently, the transcriptional activator SiwR in *M. smegmatis* was also shown to be activated in response to DNA damage (20). In *Caulobacter crescentus*, the RecA/LexA system is the main regulator of the DDR (21, 22). However, in response to DNA damage and independently of RecA/LexA, the transcriptional activator DriD directly stimulates the expression of the *didA* gene, which encodes a cell division inhibitor (12). PafB, PafC, SiwR, and DriD all belong to the widespread WYL domain-containing family of transcriptional regulators (18, 20, 23–25). In *Deinococcus* spp, DdrO, a member of the Xenobiotic Response Element (XRE) family of transcriptional regulators, is the main regulator of the DDR (15, 26–28). DdrO represses DDR genes and is proteolytically inactivated in response to DNA damage by the metalloprotease PprI (also called IrrE) (15, 29–31). Finally, in response to DNA methylation damage, the Ada-type transcriptional regulators of *Escherichia coli* and *C. crescentus* are activated post-translationally by methylation to activate the expression of genes encoding enzymes involved in repairing DNA methylation lesions (32, 33).

*Myxococcus xanthus* is found in densely populated terrestrial habitats where it is exposed to varying conditions over time and in space including exogenous genotoxic stress from factors such as desiccation, UV light, and genotoxic compounds. *M. xanthus* initiates replication of the GC-rich single-copy, circular chromosome precisely once per cell cycle, and upon completion of replication and chromosome segregation, cytokinesis follows at midcell (34–37). *M. xanthus* encodes a non-essential LexA protein, containing the conserved residues necessary for autocleavage of *E. coli* LexA and negatively autoregulates *lexA* expression (11), and two RecA proteins (38). Interestingly, transcription-based analyses previously demonstrated that the DDR in *M. xanthus* also only partially depends on LexA, suggesting that other transcription factor(s) regulate this response independently of LexA (11, 39). These transcription factor(s) remain to be identified.

Here, using a whole-cell, label-free quantitative (LFQ) proteomics approach, and mitomycin C as a DNA-damaging agent, we confirm a LexA-independent DDR *in M. xanthus*. Moreover, we identify DdiA, an XRE family transcriptional regulator, and show that it regulates part of the LexA-independent DDR. *ddiA* expression is activated heterogeneously without exogenous genotoxic stress and population-wide in response to such stress, thereby, indirectly or directly, activating the expression of *dnaE2*, which encodes the DnaE2 error-prone DNA polymerase, and inhibiting the expression of *recX*, which encodes RecX, a negative regulator of RecA. Accordingly, the Δ*ddiA* mutant has a lower mutation frequency than but also a fitness defect, suggesting that DdiA mediates a trade-off between fitness and mutagenesis. We speculate that *ddiA* expression is a tailored response to counter replication stress, thereby preventing the induction of the complete RecA/LexA-dependent DDR in the absence of exogenous genotoxic stress.

## Results

### Characterization of the proteomic response to mitomycin C treatment

Because protein abundance can be regulated at transcriptional and post-transcriptional levels, we focused on proteomic changes in response to DNA damage to map the DDR in *M. xanthus*. Specifically, we used a whole cell, LFQ proteomics approach using mitomycin C (MMC) to cause DNA damage. MMC alkylates DNA, causing interstrand crosslinks and DNA double-strand breaks (DSBs) (40). To determine the appropriate MMC concentration, we initially tested its effect on exponentially growing wild-type (WT) *M. xanthus* in suspension culture and identified 0.5 µg mL^-1^ as the highest non-lethal concentration (Fig. S1A). Therefore, to map proteome changes in response to DNA damage while minimizing cell death, MMC was added to a final concentration of 0.4 µg/mL.

Cells exposed to 0.4µg mL^-1^ MMC for 5hrs and 10hrs, corresponding to approximately one and two doubling times, continued to grow, although at a slower rate than untreated cells (Fig. 1A). At both time points, treated cells were significantly longer than untreated cells (Fig. 1B). Thus, a sublethal MMC concentration delays completion of cell division. For the LFQ proteomics analysis, total protein was extracted from four biological replicates of exponentially growing WT in suspension culture. In total 4923, 4938, and 4836 proteins were detected in untreated cells, cells treated with MMC for 5hrs and 10hrs, respectively. Using a Log_2_ (fold change (FC)) of ≥1.5 (2.83-fold increase) and ≤-1.5 (2.83-fold decrease) in protein abundance, and a -Log_10_ (*P*-value) ≥1.3 (*P*-value ≤0.05) as criteria for significant changes, 70/160 proteins and 44/196 proteins showed significantly increased and decreased abundance, respectively, in treated cells at 5/10 hrs compared to untreated cells (Fig. 1C-D; Table S1). Of these, 64 and 18 accumulated at increased and decreased levels, respectively at both time points (Fig. 1E-F). Next, we functionally categorized the differentially accumulating proteins according to the cluster of orthologous genes (COG) classification. Among the proteins with increased abundance, the largest group with a known function at both time points belonged to COG class L for proteins involved in DNA replication, recombination, and repair (14/16 at 5/10hrs with an overlap of 13) (Fig. 1E; Fig. 2; Table 1). By contrast, only a few (3/9 at 5/10hrs, respectively with an overlap of 2) COG class L proteins showed decreased abundance during MMC treatment (Fig. 1F; Fig. 2; Table 2). As expected, LexA abundance was significantly decreased at both time points of MMC treatment compared to untreated cells (Fig. 1C-D; Table 3).

**Figure 1.**
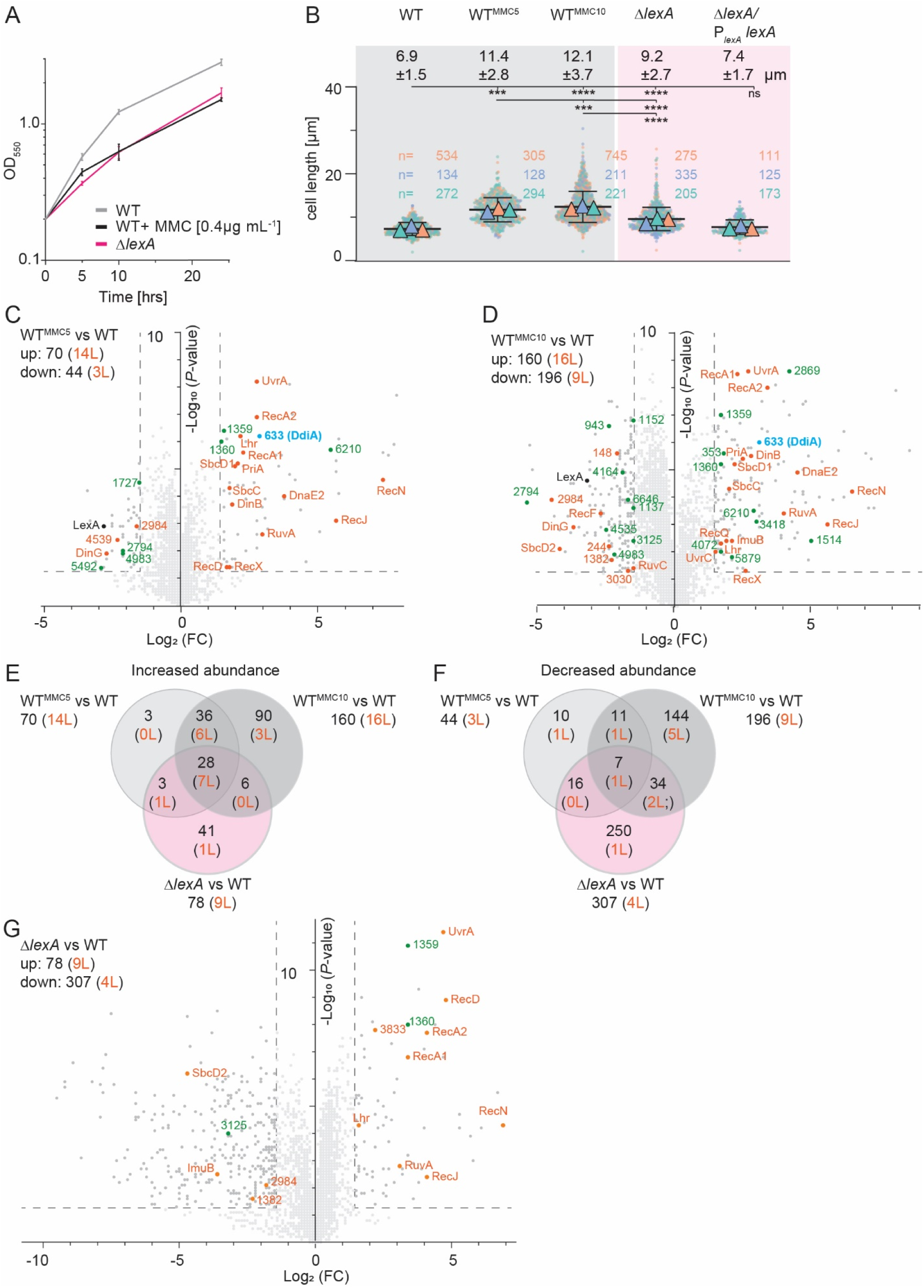
Determination of the proteomic response to MMC treatment and lack of LexA. (A) Growth curves for strains of indicated genotypes. Cells were grown in 1% CTT broth in suspension culture and MMC added as indicated. Error bars, mean ± standard deviation (STDEV) based on three independent experiments. (B) Cell length distribution of strains of indicated genotypes in the absence and presence of 0.4µg mL^-1^ MMC for 5hrs (MMC5) and 10hrs (MMC10). Measurements are included from three independent experiments indicated in different colored triangles and with the number of cells analyzed per experiment indicated in corresponding colors; error bars indicate the mean ± STDEV based on all three experiments. Numbers above indicate cell length as mean ± STDEV from all three experiments. *** *P ≤* 0.001, **** *P ≤* 0.0001, ns, not significant (2-way ANOVA multiple comparisons test). (C-D, G) Volcano plots showing differential abundance of proteins of WT treated with MMC for 5 hrs (C) and 10hrs (D) as well as the Δ*lexA* mutant relative to untreated WT (G). For all strains, samples were prepared from four biological replicates of exponentially growing cells in suspension culture. X-axis, log_2_ (FC) of proteins in experimental sample over untreated WT; Y-axis, -Log_10_ (*P*-value). Data points represent the means of four biological replicates. Significance thresholds (log_2_ (FC) of ≥1.5 (2.83-fold increase) or ≤-1.5 (2.83-fold decrease) in protein abundance and a -Log_10_ (*P*-value) ≥1.3 (*P*-value ≤0.05)) are indicated by stippled lines. Proteins of COG class L and transcriptional regulators with significantly increased or decreased abundance are indicated in orange and green, respectively; note that for the Δ*lexA* mutant, only those transcriptional regulators that had an altered abundance in MMC-treated WT are marked in green. LexA and MXAN_0633 (DdiA) are marked in black and blue, respectively. Numbers in upper, left corner indicate the total number of proteins (black) and number of COG class L proteins (orange) with significantly altered abundance. All proteins with differential abundance are listed in Table S1. (E, F) Venn diagrams of all proteins (black) and proteins of COG class L (orange) with differential abundance under the three different conditions compared to untreated WT.

**Figure 2.**
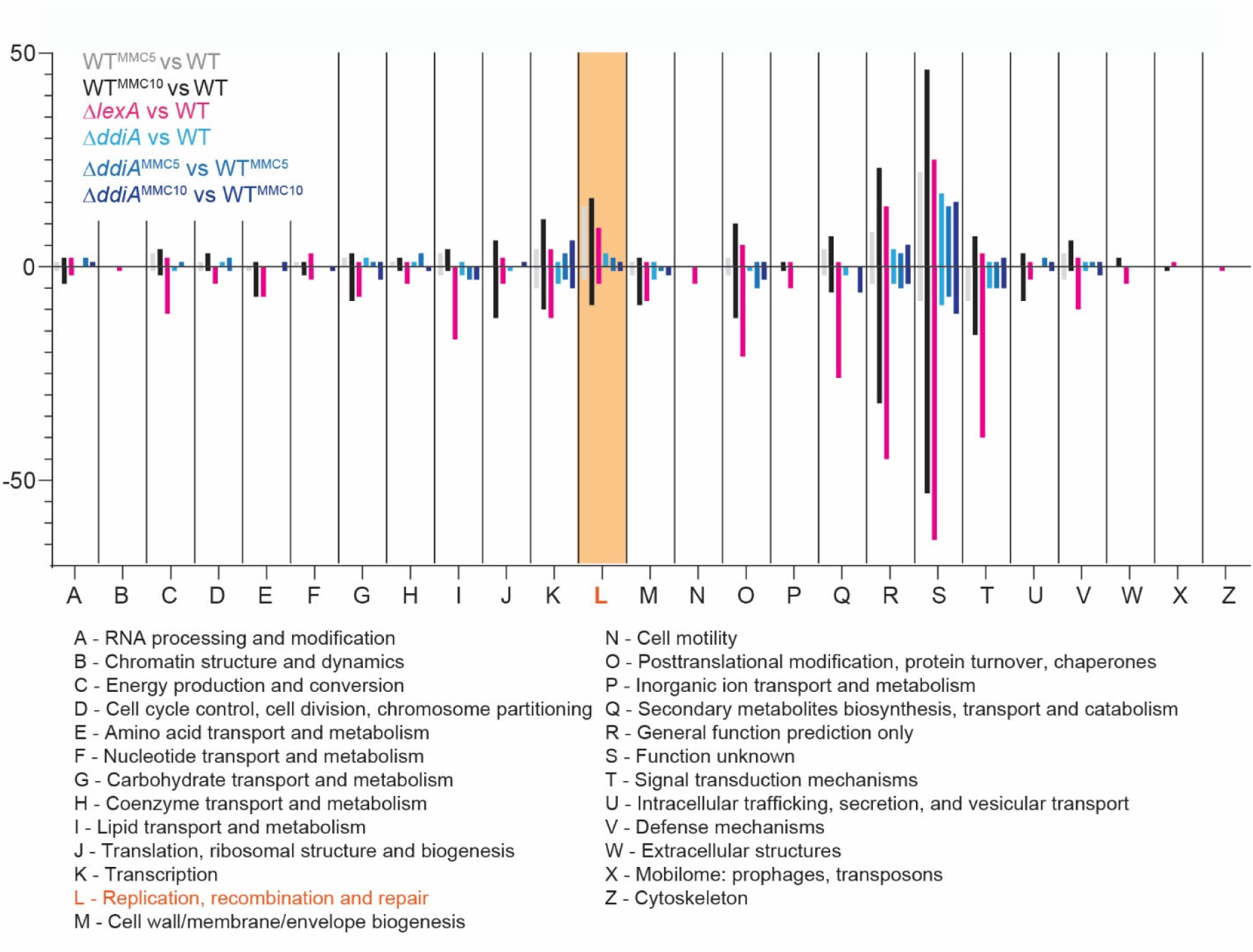
Functional classification of proteins with significantly changed abundance according to COG classes. Color code in the diagram as indicated in the upper left corner. The definition of COG classes is included below.

**Table 1.** Proteins of COG class L with increased abundance in response to MMC treatment of WT, lack of LexA, lack of DdiA, or MMC treatment of the Δ*ddiA* mutant^1, 2^.

| MXAN locus tag | Name | Protein description | WT<sup>MMC5</sup> /WT | WT<sup>MMC10</sup> /WT | $\Delta lexA$ /WT | $\Delta ddiA$ /WT | $\Delta ddiA^{MMC5}$ /WT<sup>MMC5</sup> | $\Delta ddiA^{MMC10}$ /WT<sup>MMC10</sup> |
| --- | --- | --- | --- | --- | --- | --- | --- | --- |
| 0958 | SbcD1 | SbcD subunit of SbcCD nuclease | 2.1 | 2.2 | ns | ns | ns | ns |
| 0959 | SbcC1 | SbcC subunit of SbcCD nuclease | 1.8 | 2.0 | ns | ns | ns | ns |
| 0997 | Lhr | ATP-dependent DNA helicase | 2.2 | 1.7 | 1.6 | ns | ns | ns |
| 1388 | RecA2 | Recombinase A | 2.8 | 3.4 | 4.1 | ns | ns | ns |
| 1428 | PriA | Primosomal ATP-dependent helicase | 2.0 | 2.5 | ns | ns | ns | ns |
| 1441 | RecA1 | Recombinase A | 2.3 | 2.3 | 3.4 | ns | ns | ns |
| 1651 | RuvA | RuvA subunit of RuvABC resolvase | 3.0 | 4.0 | 3.1 | ns | ns | ns |
| 1950 | RecQ | ATP-dependent DNA helicase | ns | 1.9 | ns | 1.6 | 1.7 | ns |
| 2546 | DinB | Error-prone DNA polymerase IV | 1.9 | 2.8 | ns | ns | ns | ns |
| 2609 | UvrA | UvrA subunit of UvrABC excinuclease | 2.8 | 2.7 | 4.7 | ns | ns | ns |
| 2633 | UvrC | UvrC subunit of UvrABC excinuclease | ns | 1.5 | ns | ns | ns | ns |
| 3580 | RecJ | Single-strand-specific DNA exonuclease | 5.7 | 5.6 | 4.1 | ns | ns | ns |
| 3833 |  | ATP-dependent DNA helicase | ns | ns | 2.2 | ns | ns | ns |
| 3982 | DnaE2 | Error-prone DNA polymerase E2 | 3.8 | 4.5 | ns | ns | -3.2 | -5.0 |
| 3990 | ImuB | ImuB subunit of DnaE2-ImuA-ImuB complex | ns | 2.1 | -3.6 | ns | ns | ns |
| 5350 | RecN | SMC-like DNA repair protein | 7.7 | 6.5 | 6.9 | 2.8 | ns | ns |
| 5509 | RecD | RecD subunit of RecBCD helicase/nuclease | 1.7 | ns | 4.8 | ns | ns | ns |
| 6708 | RecX | Negative regulator of RecA | 1.8 | 2.6 | ns | 1.5 | 2.9 | 2.6 |

<sup>1</sup> Numbers indicate Log<sub>2</sub> (FC) of a sample compared to the indicated control using the criteria Log<sub>2</sub> (FC) ≥ 1.5 or ≤ -1.5 and -Log<sub>10</sub> (P-value) ≥ 1.3; ns, not significant
<sup>2</sup> MMC5 and MMC10, MMC treatment for 5 and 10 hrs, respectively.

**Table 2.** Proteins of COG class L with decreased abundance in response to MMC treatment of WT, lack of LexA, lack of DdiA, or MMC treatment of the Δ*ddiA* mutant^1, 2^.

| MXAN<br>locus<br>tag | Name | Protein description | WT<sup>MMC5</sup><br>/WT | WT<sup>MMC10</sup><br>/WT | $\Delta lexA$<br>/WT | $\Delta ddiA$<br>/WT | $\Delta ddiA^{MMC5}$<br>/WT<sup>MMC5</sup> | $\Delta ddiA^{MMC10}$<br>/WT<sup>MMC10</sup> |
| --- | --- | --- | --- | --- | --- | --- | --- | --- |
| 0148 |  | RecN-like protein | ns | -2.1 | ns | ns | ns | ns |
| 0149 | SbcD2 | SbcD subunit of SbcCD nuclease complex | ns | -4.2 | -4.7 | ns | ns | ns |
| 0244 |  | AlkC-like protein for DNA alkylation repair | ns | -2.4 | ns | ns | ns | ns |
| 0246 | RecF | Component of RecFOR system | ns | -2.7 | ns | ns | ns | ns |
| 1382 |  | TatD-like exonuclease | ns | -2.3 | -2.3 | ns | ns | ns |
| 2984 |  | RecJ-like single-strand specific DNA exonuclease | -1.6 | -4.5 | -1.8 | ns | ns | ns |
| 3030 |  | MutS-related protein | ns | -1.7 | ns | ns | ns | ns |
| 4539 |  | HerA-like DNA helicase | -2.3 | ns | ns | ns | ns | ns |
| 4973 | RuvC | RuvC subunit of RuvABC resolvase | ns | -1.5 | ns | ns | ns | ns |
| 5795 | DinG | ATP-dependent DNA helicase | -2.7 | -3.7 | ns | ns | ns | ns |

<sup>1</sup> Numbers indicate Log<sub>2</sub> (FC) of a sample compared to the indicated control using the criteria Log<sub>2</sub> (FC) ≥ 1.5 or ≤ -1.5 and -Log<sub>10</sub> (P-value) ≥ 1.3; ns, not significant
<sup>2</sup> MMC5 and MMC10, MMC treatment for 5 and 10 hrs, respectively.

**Table 3.** Transcription factors with altered abundance in response to MMC treatment of WT and their abundance in the Δ*lexA* and Δ*ddiA* mutants^1, 2^.

| MXAN locus tag | Name | Protein description | WT<sup>MMC5</sup> /WT | WT<sup>MMC10</sup> /WT | $\Delta\text{lexA}$ /WT | $\Delta\text{ddiA}$ /WT | $\Delta\text{ddiA}^{\text{MMC5}}$ /WT<sup>MMC5</sup> | $\Delta\text{ddiA}^{\text{MMC10}}$ /WT<sup>MMC10</sup> |
| --- | --- | --- | --- | --- | --- | --- | --- | --- |
| 0353 |  | Sigma-54 dependent transcriptional regulator | ns | 1.8 | ns | ns | ns | ns |
| 0633 | DdiA | Transcriptional regulator, XRE family | 2.9 | 3.1 | ns | na | na | na |
| 1359 |  | Transcriptional regulator w/wHTH & WYL domains | 1.6 | 1.7 | 3.4 | ns | ns | ns |
| 1360 |  | Transcriptional regulator w/wHTH & WYL domains | 1.5 | 1.7 | 3.4 | ns | ns | ns |
| 1514 |  | ECF-sigma factor | ns | 5.0 | ns | ns | ns | ns |
| 2869 |  | Transcriptional regulator, TetR family | ns | 4.2 | ns | ns | ns | ns |
| 3418 |  | Sigma-54 dependent transcriptional regulator | ns | 3.0 | ns | ns | ns | ns |
| 4072 |  | DNA-binding response regulator, LuxR family | ns | 1.7 | ns | ns | 2.1 | ns |
| 5879 |  | Sigma-54 dependent transcriptional regulator | ns | 2.1 | ns | ns | ns | ns |
| 6210 |  | Winged helix DNA-binding domain-containing protein | 5.5 | 2.9 | ns | ns | ns | 2.5 |
| 0943 |  | Transcriptional regulator, MarR family | ns | -2.4 | ns | ns | ns | ns |
| 1137 |  | Transcriptional regulator, AraC family | ns | -1.5 | ns | ns | ns | ns |
| 1152 | RisR | Transcriptional regulator, IscR family | ns | -1.5 | ns | ns | ns | ns |
| 1727 |  | Transcriptional regulator, TetR family | -1.5 | ns | ns | ns | 1.5 | ns |
| 2794 |  | Transcriptional regulator, TetR family | -2.1 | -5.4 | ns | ns | ns | ns |
| 3125 |  | Winged helix DNA-binding domain-containing protein | ns | -1.5 | -3.2 | ns | ns | ns |
| 4164 |  | DNA-binding response regulator, OmpR_PhoB family | ns | -1.9 | ns | ns | ns | ns |
| 4446 | LexA | Regulator of SOS response | -2.8 | -3.2 | na | ns | ns | ns |
| 4535 |  | ECF-sigma factor | ns | -2.5 | ns | ns | ns | ns |
| 4983 |  | Sigma-54-dependent transcriptional regulator | -2.1 | -2.2 | ns | ns | ns | ns |
| 5492 |  | Transcriptional regulator, LysR family | -2.9 | ns | ns | ns | ns | ns |
| 6646 |  | Transcriptional regulator, MarR family | ns | -1.7 | ns | ns | ns | ns |

<sup>1</sup> Numbers indicate Log<sub>2</sub> (FC) of a sample compared to the indicated control using the criteria Log<sub>2</sub> (FC) ≥ 1.5 or ≤ -1.5 and -Log<sub>10</sub> (P-value) ≥ 1.3; ns, not significant. Note that for the $\Delta\text{lexA}$ and $\Delta\text{ddiA}$ mutants, only those transcriptional regulators that had an altered abundance in MMC-treated WT are included.
<sup>2</sup> MMC5 and MMC10, MMC treatment for 5 and 10 hrs, respectively.

The COG class L proteins with increased abundance upon MMC exposure included proteins for homologous recombination (HR) and DSB repair (RecA1, RecA2, RecD, RecN, the SbcC1D1 proteins, and RuvA) as well as nucleotide excision repair (NER) (UvrA and UvrC) (Table 1). Moreover, proteins for error-prone DNA repair (DnaE2, ImuB, and DinB) had increased abundance. Also, seven helicases and nucleases were more abundant. Finally, RecX, the negative regulator of RecA that inhibits RecA recombinase activity and coprotease activity (2, 41), was more abundant at both time points. The COG class L proteins with decreased abundance (Table 2) included proteins involved in HR and DSB (SbcD2, RecF, RuvC), various helicases and nucleases as well as proteins possibly involved in the repair of alkylated DNA (MXAN_0244), and mismatch repair (MXAN_3030).

Consistent with these findings, Campoy et al. (11) reported that *recA2* and *recN* expression was strongly upregulated in response to 8hrs of MMC treatment. They also found that *ssb* expression was strongly upregulated by MMC treatment, while *recA1* and *ruvA* expression was unaffected. We observed that Ssb accumulated at similar levels in untreated and MMC-treated cells at 5 and 10hrs and that RecA1 and RuvA accumulated at increased levels at both time points. However, we also note that in (11), cells were treated with 40µg/mL MMC, which caused cell death under the conditions we used (Fig. S1A), making direct comparisons difficult.

### Characterization of the LexA-dependent proteomic DNA damage response

Next, we asked which of the proteomic changes in response to MMC treatment were regulated by LexA. Among the two RecA proteins in *M. xanthus*, Norioka et al. (38) reported that RecA1 is not essential, while RecA2 may be essential. Sheng et al. (42) found that the two *recA* genes could be inactivated individually, but a double mutant lacking both RecA proteins could not be obtained. It was previously reported that LexA is non-essential in *M. xanthus* (11, 39). Therefore, to map the LexA-dependent proteomic DDR, we generated an in-frame *lexA* deletion mutant (Δ*lexA*) in which the first 10 codons of the 223-codon *lexA* gene were fused in-frame to the last 10 codons. We readily obtained Δ*lexA* mutants. Because LexA has been reported to be essential or, alternatively, that lack of LexA causes strong growth defects in other bacteria (21, 43–46), we characterized three independent Δ*lexA* mutants. The three Δ*lexA* mutants behaved similarly and had a slight growth defect, consistent with previous reports (39), and the growth rate was similar to that of WT treated with 0.4µg mL^-1^ MMC (Fig. 1A). Unlike previously reported (39), we observed that the Δ*lexA* cells were significantly longer than untreated WT cells but shorter than MMC-treated WT cells (Fig. 1B). The cell length defect was corrected by the ectopic expression of *lexA* from its native promoter (P*_lexA_*) on a plasmid integrated in a single copy at the Mx8 *attB* site (Fig. 1B; Fig. S1B). We speculate that the difference between the previously published Δ*lexA* mutant (39) and our mutant could be caused by the use of different in-frame deletion mutants as well as differences in growth media and temperature.

Next, we determined the proteome in four biological replicates of exponentially growing suspension cultures of the Δ*lexA* mutant. The LFQ proteomics analysis detected a total of 4812 proteins in the Δ*lexA* mutant (Fig. 1G). Compared to untreated WT, 78 proteins were increased and 307 proteins decreased in abundance (Fig. 1G; Table S1). Of the 78 proteins with increased abundance, 37 were also upregulated in one or both samples of MMC-treated WT cells (Fig. 1E). Of the 307 proteins with decreased abundance, 57 were also less abundant in one or both samples of MMC-treated WT cells (Fig. 1F).

Among the proteins with increased abundance in the Δ*lexA* mutant, the largest group with a known function belonged to COG class L (Fig. 2). Specifically, nine COG class L proteins were more abundant in the Δ*lexA* mutant, eight of which overlapped with COG class L proteins that were also more abundant in MMC-treated WT. These included proteins involved in HR, DSB repair, and NER (Fig. 1E; Table 1). The one protein (MXAN_3833) that accumulated at an increased level only in the Δ*lexA* mutant is an ATP-dependent helicase with an ill-defined function in DNA repair (Table 1). Notably, nine COG class L proteins that displayed increased abundance in MMC-treated WT accumulated at unchanged or even lower levels (ImuB) in the Δ*lexA* mutant, including proteins involved in HR, DSB repair, NER, and all three proteins involved in error-prone DNA repair, as well as RecX (Fig. 1E; Table 1). Among the proteins with decreased abundance in the Δ*lexA* mutant, the largest group with a known function belonged to COG class T for signal transduction mechanisms (Fig. 2).

Among the four COG class L proteins with decreased abundance, three overlapped with proteins that also had decreased abundance in MMC-treated WT cells, and the one protein that was only downregulated in the Δ*lexA* mutant was ImuB of the DnaE2/ImuA/ImuB translesion DNA polymerase complex (Fig. 1F; Table 1-2). Among the COG class L proteins with decreased abundance in MMC-treated WT cells, seven were not downregulated in the Δ*lexA* mutant (Fig. 1F; Table 2). Thus, LexA regulates the abundance of only some of the proteins involved in the MMC-induced DDR, while other such proteins are regulated independently of LexA.

In agreement with our proteomics-based observations, *recA2*, *uvrA*, and *lhr* expression were reported to be negatively regulated by LexA, and *sbcC1*, *sbcD1,* and *dnaE2* upregulation in response to UV-induced DNA damage was reported to be LexA-independent (11, 39). It was also reported that *recA1* and *ruvA* expression is LexA-independent (11), while *recN* expression was reported to be either LexA-independent (11) or LexA-dependent (39). Thus, there is partial agreement between the transcriptomic- and proteomic-based analyses of the Δ*lexA* mutant. However, we note that the Δ*lexA* mutants used by (11, 39) and our Δ*lexA* mutant are different and cells were grown in different growth media and at different temperatures, making direct comparisons between the transcription-based and proteomics results difficult. Nevertheless, both the transcription-based data (11, 39) and our proteomics data document that the DDR includes a LexA-dependent and a LexA-independent response. This conclusion is further supported by the observation that the Δ*lexA* mutant has a cell length phenotype distinct from that of WT cells treated with a sublethal MMC concentration (Fig. 1B).

### Identification of DdiA, an XRE family transcriptional regulator upregulated by MMC treatment independently of LexA

The partial agreement between the transcriptomic- and proteomic-based analyses of the Δ*lexA* mutant supports that at least part of the LexA-independent DDR in *M. xanthus* is regulated at the transcriptional level. To identify transcriptional regulators potentially involved in this response, we first identified those with altered abundance in MMC-treated WT. Subsequently, we identified those that did not change in abundance in the absence of LexA.

In addition to LexA, we identified 10 and 11 transcriptional regulators with increased and decreased abundance, respectively in MMC-treated WT (Fig. 1C-D; Table 3). Among these, three had LexA-dependent changes in abundance (Fig. 1G; Table 3). Interestingly, two of these (MXAN_1359 and _1360) are homologs of PafB, PafC, SiwR, and DriD that regulate LexA-independent DDRs in *M. smegmatis*, *M. tuberculosis* and *C. crescentus*. Nevertheless, because LexA regulates MXAN_1359 and _1360 abundance, these proteins were not investigated further. Most of the COG class L proteins with increased abundance in response to MMC treatment and not regulated by LexA, accumulated at higher levels at both 5 and 10hrs (Fig. 1E; Table 1). Therefore, we next focused on those transcriptional regulators with altered abundance at both time points of MMC treatment. Among these, two candidates (MXAN_0633 and _6210) were upregulated, and two (MXAN_2794 and _4983) were downregulated at both time points (Fig. 1C-D; Table 3). Of these four transcriptional regulators, only MXAN_0633, which accumulated at highly increased levels in response to MMC treatment, belongs to a transcription factor family involved in a LexA-independent DDR. Consequently, from here on, we focused on MXAN_0633, which we refer to as <u>D</u>NA <u>d</u>amage-induced protein <u>A</u> (DdiA).

Based on sequence analysis, DdiA, similar to DdrO of *Deinococcus* spp., belongs to the XRE family of transcriptional regulators and has an N-terminal helix-turn-helix (HTH) DNA binding domain of the XRE-type and a C-terminal domain predicted to mediate oligomerization. Indeed, a high-confidence AlphaFold-Multimer structural model of DdiA supports that DdiA forms a dimer with an N-terminal HTH domain and two C-terminal α-helices mediating dimerization (Fig. 3A; Fig. S2A). *ddiA* is located adjacent to the *pomXYZ* genes, which encode the cell division regulators PomX, PomY and PomZ (36, 37), the *ddiA* locus is conserved in closely related myxobacteria (Fig. S2B), and *ddiA* is not predicted to be part of an operon (47) (Fig. S2C).

**Figure 3.**
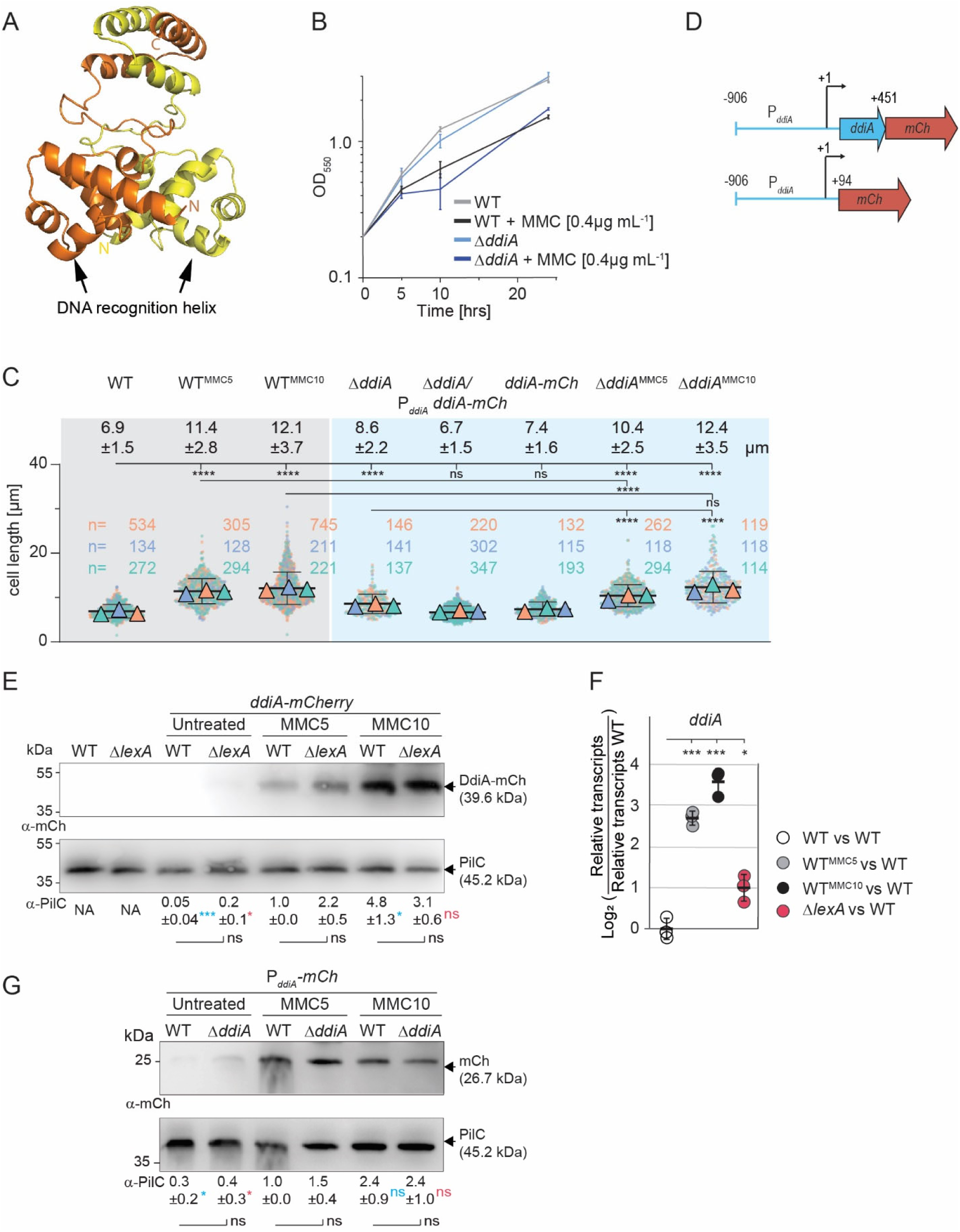
*ddiA* expression and DdiA abundance are induced by MMC treatment independently of LexA and DdiA. (A) AlphaFold-Multimer structural model of DdiA dimer. Protomers are shown in yellow and orange with the N-termini indicated in the same color. Model rank 1 is shown. (B) Growth of strains of indicated genotypes. Cells were grown in 1% CTT broth in suspension culture and MMC added as indicated. Error bars, mean ± STDEV based on three independent experiments. Note that the growth curves of WT with or without MMC are the same as in Fig. 1A. (C) Lack of DdiA causes a cell length defect. Cells were analyzed as described in the legend to Fig. 1B. Note that the WT samples are the same as in Fig. 1. **** *P ≤* 0.0001, ns, not significant (2-way ANOVA multiple comparisons test). (D) Schematic of the construct for the P*_ddiA_*-DdiA-mCh protein fusion (upper) and the P*ddiA*-mCh promoter fusion (lower). +1 indicates the transcription start site of *ddiA,* +94 and +451 indicate the first nucleotide of the start codon and the last nucleotide of the last coding codon in *ddiA*, respectively. (E) Immunoblot analysis of DdiA-mCh abundance. Cells were harvested from exponentially growing cells in suspension culture and protein from 7×10^5^ cells per sample loaded; the PilC blot served as a loading control. Samples marked MMC5 and MMC10 are from cells grown in the presence of MMC for 5 and 10hrs, respectively. For quantification, DdiA-mCh signals were corrected relative to the PilC loading control and normalized relative to the WT MMC5 sample, which was set to 1.0. Numbers below indicate intensity of the DdiA-mCh signal as mean ± STDEV based on three biological replicates; NA, not applicable. * *P* ≤ 0.05, ** *P* ≤ 0.01, *** *P* ≤ 0.001, ns not significant (Unpaired t-test with Welch’s correction). Samples marked with an asterisk in blue were compared to the WT MMC5 sample, an asterisk marked in red to the Δ*lexA* mutant MMC5 sample, and in black to the sample from the same time point. (F) RT-qPCR analysis of *ddiA* transcript levels. Total RNA was isolated from exponentially growing cells in suspension culture and in the presence of MMC as indicated. Data are shown as log_2_ of transcript in a strain relative to that of the untreated WT. Individual data points represent three biological replicates with each two technical replicates and are colored according to the strain analyzed. Error bars indicate mean ± STDEV. * *P* ≤ 0.05, *** *P* ≤ 0.001 (Unpaired t-test with Welch’s correction). (G) Immunoblot analysis of P*_ddiA_* expression. P*_ddiA_* was fused to a promoterless *mCh* (see panel D). Cells were grown, treated, and analyzed as in panel (E). For quantification, the mCh signals were corrected relative to the PilC loading control and normalized relative to the WT MMC5 sample, which was set to 1.0. Numbers below indicate intensity of the mCh signal as mean ± STDEV based on three biological replicates; * *P* ≤ 0.05, ns not significant (Unpaired t-test with Welch’s correction). Samples marked in blue were compared to the WT MMC5 sample, in red to the Δ*ddiA* mutant MMC5 sample, and in black to the sample from the same time point.

### MMC treatment induces *ddiA* transcription and DdiA abundance independently of LexA

To investigate the potential role of DdiA in the MMC-induced DDR, we generated an in-frame *ddiA* deletion mutant (Δ*ddiA*) in which the first 10 codons of the 120-codon *ddiA* gene were fused in-frame to the last 10 codons. The Δ*ddiA* mutant had a growth rate similar to WT (Fig. 3B; but see also below). However, cells of the Δ*ddiA* mutant were significantly longer than WT cells (Fig. 3C). This cell length defect was corrected by the ectopic expression of *ddiA-mCherry* (*ddiA-mCh*) from the native *ddiA* promoter (P*_ddiA_*) on a plasmid integrated in a single copy at the Mx8 *attB* site (Fig. 3C-D). Furthermore, when *ddiA-mCh* was expressed from the *ddiA* locus, these cells were indistinguishable from WT cells, demonstrating that the DdiA-mCh fusion is fully active.

To corroborate that MMC treatment increases DdiA abundance independently of LexA, we took advantage of the DdiA-mCh fusion expressed from the *ddiA* locus. In immunoblots, DdiA-mCh was barely detected in untreated WT and Δ*lexA* cells but was strongly upregulated at both 5 and 10hrs of MMC treatment in both strains (Fig. 3E; Fig. S3A). Importantly, DdiA-mCh abundance was similar in WT and the Δ*lexA* mutant in both untreated and MMC-treated cells (Fig. 3E). To determine at which level DdiA abundance is regulated in response to MMC treatment, we performed RT-qPCR analyses. In WT, the *ddiA* transcript level was significantly higher at 5 and 10hrs of MMC treatment compared to untreated cells (Fig. 3F). In the Δ*lexA* mutant, the *ddiA* transcript level was slightly, although significantly, higher compared to the untreated WT but still significantly lower than in MMC-treated WT (Fig. 3F). To determine whether DdiA autoregulates *ddiA* transcription, we generated a P*_ddiA_*-*mCh* promoter fusion in which P*_ddiA_* was fused to *mCh* (Fig. 3D) on a plasmid integrated in a single copy at the Mx8 *attB* site in the WT and the Δ*ddiA* mutant. In agreement with the RT-qPCR analysis, mCh abundance increased significantly in MMC-treated WT as assessed by immunoblotting; mCh abundance also increased significantly in the MMC-treated Δ*ddiA* mutant; importantly, mCh abundance was similar in WT and the Δ*ddiA* mutant in untreated cells as well as in cells treated with MMC for 5 and 10hrs (Fig. 3G; Fig. S3B).

Taken together, these results demonstrate that MMC treatment induces *ddiA* transcription, leading to increased DdiA abundance independently of LexA, and DdiA neither positively nor negatively autoregulates *ddiA* expression.

### DdiA activates *dnaE2* and represses *recX* transcription in response to MMC treatment

The Δ*ddiA* mutant, similar to WT, responded with a slightly reduced growth rate to 0.4µg mL^-1^ MMC (Fig. 3B). As in WT, the Δ*ddiA* cells exhibited an increased cell length in response to MMC treatment for 5 and 10hrs (Fig. 3C). In the LFQ proteomics experiments with the Δ*ddiA* mutant, a total of 4315 (untreated), 4311 (5hrs MMC), and 4318 (10hrs MMC) proteins were detected (Fig. 4A-C). In comparison to untreated WT, untreated Δ*ddiA* cells had an increased abundance of 33 proteins, including three COG class L proteins (RecN, RecX, RecQ), and a decreased abundance of 33 proteins, none of which belonged to COG class L (Fig. 4A; Table 1-2). At 5hrs of MMC treatment, the abundance of 33 proteins, including two COG class L proteins (RecX, RecQ), was significantly increased, and 32 proteins, including one COG class L protein (DnaE2), were significantly decreased compared to WT treated with MMC for 5hrs (Fig. 4B; Table 1-2). At 10hrs of MMC treatment, the abundance of 36 proteins, including one COG class L protein (RecX), was significantly increased, and 49 proteins, including one COG L class protein (DnaE2), were significantly decreased compared to WT treated with MMC for 10hrs (Fig. 4C; Table 1-2). At all time points, LexA accumulated in the Δ*ddiA* mutant as in the WT (Table 3). Among the four COG class L proteins (RecN, RecQ, RecX, and DnaE2) affected by the lack of DdiA, only RecN is regulated by LexA (Table 1). The increased RecN abundance in untreated Δ*ddiA* cells suggests that DdiA may inhibit RecN accumulation in these cells. However, the increased RecN abundance in response to MMC treatment in WT can be explained by LexA regulation, and during MMC treatment, DdiA does not significantly regulate RecN abundance (Table 1).

**Figure 4.**
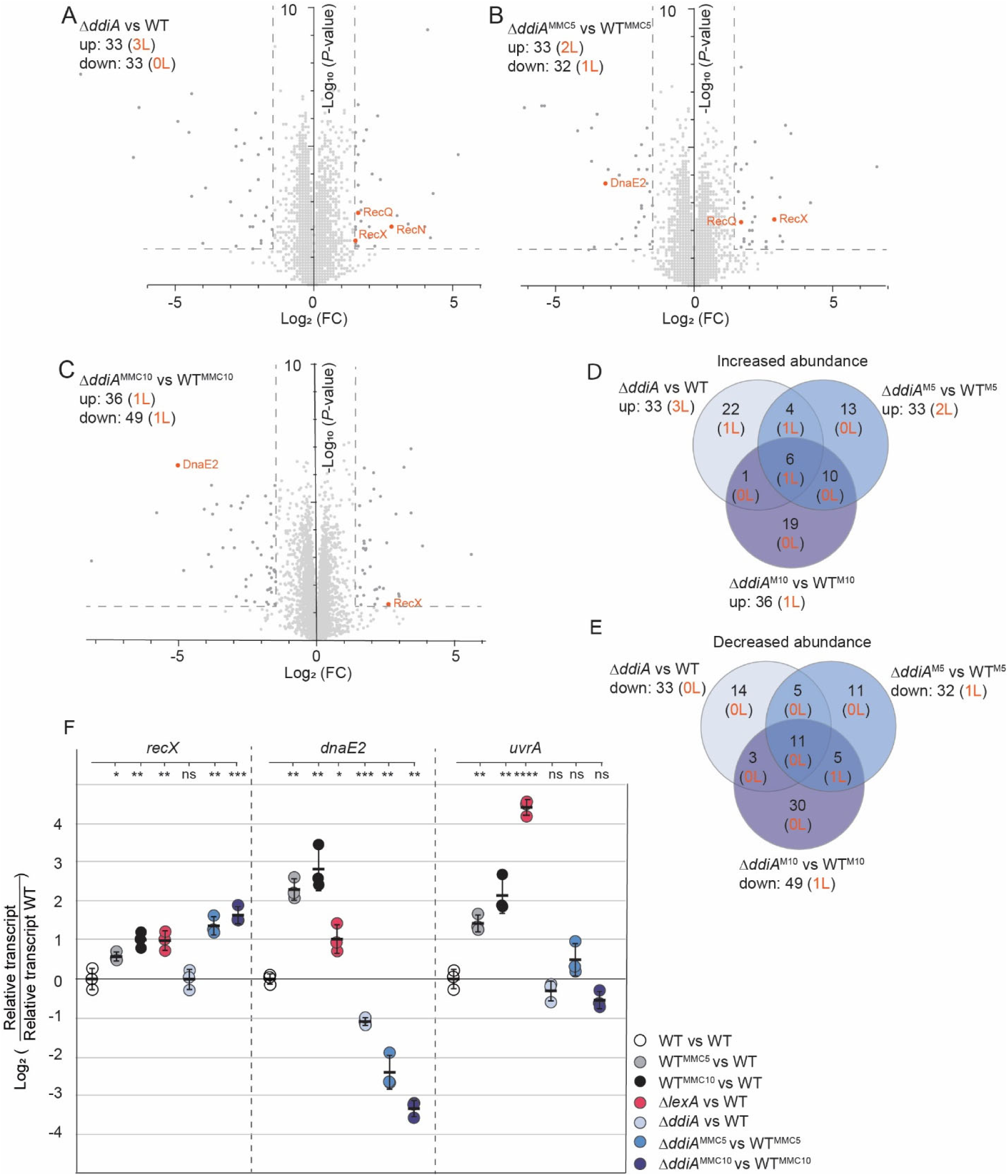
DdiA activates *dnaE2* and represses *recX* transcription in response to MMC treatment. (A-C) Determination of the proteomic response to lack of DdiA and MMC treatment of the Δ*ddiA* mutant. Volcano plots showing differential abundance of proteins of the untreated Δ*ddiA* mutant relative to untreated WT (A), the Δ*ddiA* mutant treated with MMC for 5 hrs (B) and 10 hrs (C) relative to WT treated with MMC for 5 and 10hrs, respectively. Samples were prepared from four biological replicates. Strains were grown as in Fig. 1C. Data points represent means of four biological replicates. Significance thresholds as in Fig. 1C and indicated by stippled lines. Proteins of COG classes L with significantly increased or decreased abundance are indicated in orange. Numbers in upper, left corner indicate the total number of proteins and the number of COG class L proteins (orange) with significantly altered abundance. All proteins with differential abundance are listed in Table S1. (D, E) Venn diagrams of all proteins (black) and proteins of COG class L (orange) with differential abundance under the three different conditions relative to WT. (F) RT-qPCR analysis of *recX, dnaE2,* and *uvrA* transcript levels. Cells were grown as in Fig. 3F. Data are shown as Log_2_ of transcript in a strain relative to that of the untreated WT or WT treated with MMC for 5 or 10hrs. Individual data points represent three biological replicates with each two technical replicates and are colored according to the strain analyzed. Error bars indicate mean ± STDEV. * *P* ≤ 0.05, ** *P* ≤ 0.01, *** *P* ≤ 0.001, **** *P* ≤ 0.0001, ns not significant (Unpaired t-test with Welch’s correction).

To determine at which level DdiA affects the abundance of RecQ, RecX, and DnaE2, we focused on RecX and DnaE2 because they accumulated at increased and decreased levels, respectively in the Δ*ddiA* mutant at both time points of MMC treatment compared to WT (Table 1). Using RT-qPCR, we observed that *recX* transcription was slightly but significantly induced in MMC-treated WT (Fig. 4F) as well as in the Δ*lexA* mutant compared to the untreated WT, but not in the untreated Δ*ddiA* mutant (Fig. 4F). Importantly, *recX* transcription was significantly higher in MMC-treated Δ*ddiA* cells compared to MMC-treated WT (Fig. 4F). *dnaE2* transcription was highly induced in MMC-treated WT and only slightly, although significantly, induced in the Δ*lexA* mutant (Fig. 4F). By contrast, *dnaE2* expression was significantly lower in the untreated Δ*ddiA* mutant compared to WT as well as in the MMC-treated Δ*ddiA* mutant compared to MMC-treated WT at both time points (Fig. 4F). As a control, we focused on UvrA that increased in abundance in MMC-treated WT as well as in the Δ*lexA* mutant independently of DdiA (Table 1). Transcription of *uvrA* was strongly induced in MMC-treated WT at both time points and in the Δ*lexA* mutant (Fig. 4F). By contrast, *uvrA* expression in the Δ*ddiA* mutant was as in WT under all three conditions (Fig. 4F).

We conclude that DdiA, directly or indirectly, represses transcription of *recX* during MMC treatment, and activates transcription of *dnaE2* in untreated as well as in MMC-treated cells.

### *ddiA* expression is activated heterogeneously in the absence of exogenous genotoxic stress

Intriguingly, we serendipitously observed by fluorescence microscopy that a subpopulation (10.1±1.4%) of the cells that synthesizes the fully active DdiA-mCh fusion from the native locus accumulated DdiA-mCh when grown under standard conditions without exogenous genotoxic stress (Fig. 5A). In these cells, DdiA-mCh co-localized perfectly with the 4,6-diamidino-2-phenylindole (DAPI)-stained nucleoid, strongly supporting that DdiA is a DNA-binding protein (Fig. 5A). DdiA-mCh^+^ cells varied in length but were overall significantly longer than DdiA-mCh^-^ cells (Fig. 5A). Under the same conditions, the P*_ddiA_*-*mCh* promoter fusion was also heterogeneously expressed in WT and in the Δ*ddiA* mutant (Fig. 5B). In both strains, the mCh^+^ cells were of varying lengths but overall significantly longer than the mCh^-^ cells (Fig. 5B). Notably, the mCh^+^ cells of the Δ*ddiA* mutant were significantly longer than the mCh^+^ cells of the WT (Fig. 5B). The translational *ddiA-mCh* fusion expressed from the native site was also heterogeneously expressed in the Δ*lexA* mutant (Fig. 5C).

**Figure 5.**
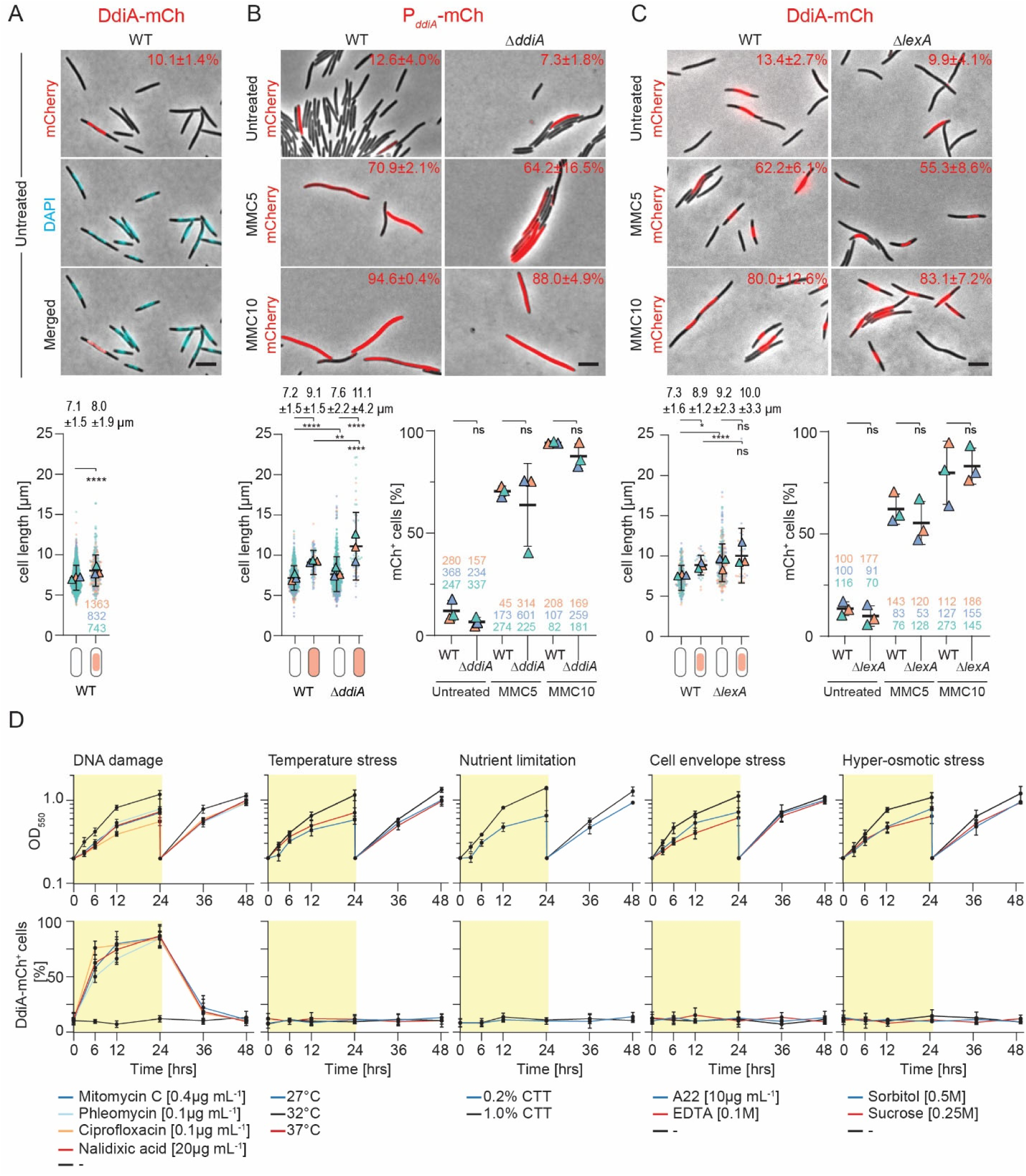
*ddiA* expression is reversibly activated by genotoxic stress. (A) *ddiA* expression is activated heterogeneously in the absence of exogenous genotoxic stress. Upper panels, untreated WT cells expressing DdiA-mCh from the native site and stained with DAPI were visualized by fluorescence microscopy and phase-contrast microscopy. Phase-contrast and fluorescence images of representative cells were merged. Number in upper right indicates fraction of DdiA-mCh^+^ cells based on three biological replicates. Lower panel, cell length distribution of DdiA-mCh^-^ and DdiA-mCh^+^ cells. Measurements are included from the three biological replicates indicated in differently colored triangles and with the number of cells analyzed per experiment indicated in corresponding colors; error bars, mean ± STDEV based on all three experiments. Numbers above indicate cell length as mean ± STDEV from all three experiments. **** *P* ≤ 0.0001 (2-way ANOVA multiple comparisons test). Scale bar, 5 mm. (B, C) *ddiA* expression is reversibly activated population-wide by MMC independently of DdiA and LexA. Cells expressing the P*_ddiA_*-*mCh* promoter fusion (B) and the DdiA-mCh protein fusion (C) in the absence (untreated) and presence of MMC for 5 (MMC5) and 10hrs (MMC10) were visualized as in (A). Merged images of representative cells are shown. Numbers in upper right corners indicate the fraction of mCh^+^/DdiA-mCh^+^ cells based on three biological replicates. Lower panel left: cell length distribution of DdiA-mCh^-^ and DdiA-mCh^+^ cells presented as in (A). Numbers above indicate cell length as mean ± STDEV from all three experiments. * *P ≤* 0.05, ** *P ≤* 0.01 **** *P ≤* 0.0001, ns not significant (2-way ANOVA multiple comparisons test). Scale bars, 5 mm. Lower panel right: Comparison of the fraction of mCh^+^/DdiA-mCh^+^ cells in the indicated strains in the absence and presence of MMC. ns, not significant (Unpaired t-test with Welch’s correction). Numbers below indicate the number of cells analyzed in the three biological replicates. (D) *ddiA* expression is reversibly activated by DNA damage. WT cells expressing DdiA-mCh from the native site were exposed to the indicated stressors, growth followed (upper panels) and the fraction of DdiA-mCh^+^ cells quantified (lower panels). Exponentially growing cells in suspension culture were exposed to a specific stress at 0hrs, at time point 24hrs, the stressor was removed and cultures diluted; samples were withdrawn at the indicated time points and analyzed by fluorescence microscopy. Upper and lower panels, error bars, mean ± STDEV from three biological replicates; at each time point, > 100 cells were analyzed by fluorescence microscopy per replicate.

Upon MMC treatment for 5 and10hrs, the fraction of mCh^+^ cells in WT and the Δ*ddiA* mutant expressing the P*_ddiA_*-*mCh* promoter fusion dramatically increased (Fig. 5B). Similarly, the fraction of DdiA-mCh^+^ cells in WT and in the Δ*lexA* mutant expressing the translational *ddiA-mCh* fusion from the native locus dramatically increased in response to MMC treatment (Fig. 5C).

Altogether, we conclude that (1) P*_ddiA_* is activated heterogeneously in the absence of exogenous genotoxic stress, and this activation is independent of DdiA and LexA. (2) The low DdiA-mCh levels in the absence of exogenous genotoxic stress in the population-based immunoblot analyses (Fig. 3E) mask that *ddiA-mCh* is activated heterogeneously, with a small fraction of cells expressing *ddiA-mCh* and accumulating DdiA-mCh. (3) MMC treatment induces *ddiA* transcription and DdiA-mCh accumulation independently of LexA and DdiA, in agreement with the population-based immunoblot analyses and RT-qPCR analyses (Fig. 3E-G). (4) The MMC-induced *ddiA-mCh* transcription is not restricted to the “original” mCh^+^ cells but occurs throughout the population. Finally, the observations that (1) the DdiA-mCh^+^/mCh^+^ cells in the WT are generally longer than the DdiA-mCh^-^/mCh^-^ cells, and (2) the mCh^+^ cells in the Δ*ddiA* mutant are longer than mCh^+^ cells in the WT, suggest that the cue inducing *ddiA* expression results in a cell length defect and lack of DdiA exacerbates this phenotype.

### *ddiA* expression is reversibly induced by DNA damage

To examine whether the increased *ddiA* expression is a specific response to the DNA damage caused by MMC, a more general response to DNA damage, or a response to general cellular stress, we exposed WT cells expressing *ddiA-mCh* from the native *ddiA* locus to various DNA damaging agents and other stresses. Subsequently, we tracked DdiA-mCh induction at the single-cell level using fluorescence microscopy. As observed for a sublethal concentration of MMC, sublethal concentrations of ciprofloxacin (0.1µg mL^-1^) and nalidixic acid (20µg mL^-1^), both of which inhibit topoisomerase IV and DNA gyrase (48), resulting in protein-linked DNA breaks, DSBs and inhibition of DNA replication, caused a dramatic increase in the fraction of DdiA-mCh^+^ cells over 24hrs of exposure (Fig. 5D; Fig. S4). Similarly, exposure to a sublethal concentration of phleomycin (0.1µg mL^-1^), which binds DNA and directly induces DNA breaks and DSBs (49), caused a dramatic increase in the fraction of DdiA-mCh^+^ cells over 24hrs of exposure (Fig. 5D; Fig. S4). Furthermore, 12-24hrs after removal of these DNA-damaging compounds, the fraction of DdiA-mCh^+^ cells had returned to the pre-treatment level (Fig. 5D). By contrast, exposure to stress conditions such as growth at decreased (27°C) or increased (37°C) temperatures, low nutrient levels (0.2% casitone), the highest sublethal concentration of EDTA (100 µM), which disrupts outer membrane integrity, A22 (10 µg mL^-1^), which interferes with peptidoglycan biosynthesis (50), or hyper-osmotic stress did not lead to an increase in the fraction of mCh^+^ cells (Fig. 5D; Fig. S4).

We conclude that *ddiA-mCh* expression is reversibly activated by DNA damage and returns to the pre-treatment pattern upon removal of the stressor.

### Lack of DdiA results in a reduced mutation frequency but also a fitness defect

Error-prone DNA repair significantly contributes to DNA damage-induced mutagenesis (4, 7). Given that DdiA, directly or indirectly, activates the expression of *dnaE2*, we hypothesized that the Δ*ddiA* mutant would have a lower mutation frequency than WT in the absence of exogenous genotoxic stress. To test this, we used a rifampicin resistance (Rif^R^) assay, in which point mutations in the *rpoB* gene, which encodes the β-subunit of the RNA polymerase, can be detected because they confer Rif^R^ (51, 52). We grew WT and the Δ*ddiA* mutant under standard conditions in suspension culture, plated cells on standard solid growth medium containing 25µg/mL Rif, and then counted the number of Rif^R^ colony forming units. While the WT had a mutation frequency of ∼40 per 2×10^8^ cells, the Δ*ddiA* mutant had a 4-5-fold lower frequency (Fig. 6A). This fold-reduction is similar to those reported for *dnaE2*, *imuA,* and *imuB* mutants in *M. xanthus* in the absence of exogenous genotoxic stress (53, 54).

**Figure 6.**
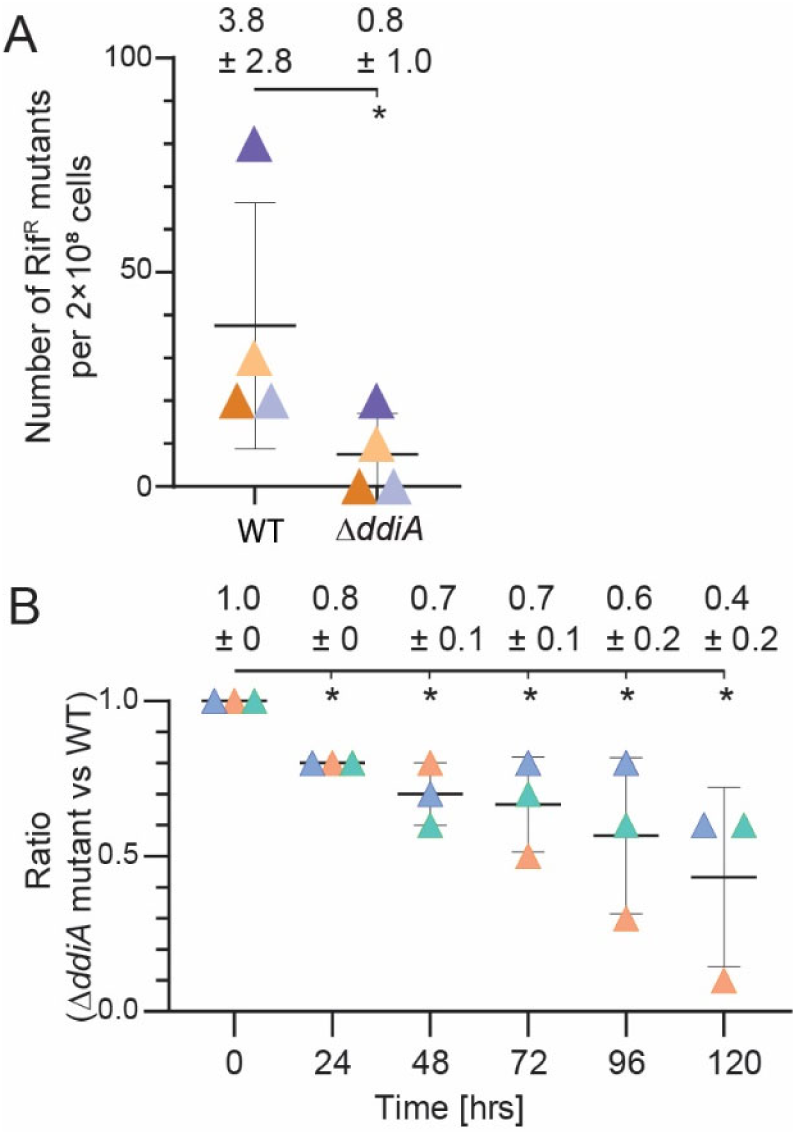
Lack of DdiA results in a reduced mutation frequency but also a fitness defect in the absence of exogenous genotoxic stress. (A) Lack of DdiA results in a reduced mutation frequency in the absence of exogenous genotoxic stress. Cultures of the indicated exponentially growing strains were plated on standard growth medium without (for viable cell counts) and with 25 μg/mL rifampicin to score Rif^R^ CFU. Mutation frequencies were calculated by dividing the numbers of Rif^R^ CFU by the number of cells analyzed in the experiments. Numbers above, mutation frequency based on four biological replicates indicated in different colors. Error bars, mean ± STDEV from four biological replicates. * *P ≤* 0.05 (Paired t-test). (B) Lack of DdiA results in a fitness defect in the absence of exogenous genotoxic stress. WT and the Δ*ddiA* mutant were mixed in suspension culture at a 1:1 ratio and growth of the mixed culture followed by measuring OD_550_. Cells were kept in the exponential growth phase in suspension culture by repeated dilution. The ratio of the WT to the Δ*ddiA* mutant was determined using a qPCR approach (Materials and Methods). Error bars, mean ± STDEV based on three biological replicates indicated in different colors. * *P ≤* 0.05 (Unpaired t-test).

When grown separately, the Δ*ddiA* mutant had a growth rate similar to WT (Fig. 3B). To assess the potential subtle fitness effects of the Δ*ddiA* mutation, we used a more sensitive competition growth experiment, in which the Δ*ddiA* mutant and WT were grown in co-culture. The ratio of the Δ*ddiA* mutant to WT was measured using a quantitative PCR (qPCR) approach immediately after mixing the two cultures and followed for several days of growth in suspension culture under standard conditions in which cells were kept in the exponential growth phase by repeated dilution. Starting from a 1:1 ratio of the Δ*ddiA* mutant to WT, WT consistently outcompeted the Δ*ddiA* mutant (Fig. 6B). Thus, DdiA provides a fitness advantage in the absence of exogenous genotoxic stress.

## Discussion

Here, we confirm the existence of a DDR in *M. xanthus* and that LexA is non-essential and involved in regulating this response (11, 39). We also confirm the previous transcriptomic-based suggestion that only part of the DDR is regulated by LexA and that other transcription factor(s) are involved in regulating the DDR in *M. xanthus* (11, 39). Using a candidate approach, we identify the transcriptional regulator DdiA and show that it regulates part of the LexA-independent DDR.

Using MMC as a DNA damaging agent, we found that the DDR at the proteomic level includes proteins involved in HR, DSB repair, NER, error-prone DNA repair, and RecX, the negative regulator of RecA. Among these, several accumulated at unchanged levels or even at a lower level in the Δ*lexA* mutant and, therefore, were categorized as LexA-independent. These proteins include proteins involved in HR, DSB repair, and NER, all three proteins involved in error-prone DNA repair (ImuB, DnaE2, DinB) that were upregulated during the MMC-induced DDR, as well as RecX.

DdiA is a member of the XRE family of transcriptional regulators. This family is abundant and widespread in bacteria (55) and functions as both transcriptional repressors and activators (56). In the absence of DdiA, the abundance of 154 proteins was significantly altered in untreated and/or MMC-treated cells, with only some of these proteins involved in the DDR. Because *ddiA* expression is induced by different types of DNA damage and not by general cellular stress, we discuss DdiA in the context of DNA damage. Without exogenous genotoxic stress, *ddiA* is expressed and DdiA accumulates independently of LexA and DdiA in a subpopulation of cells. In response to DNA damage, *ddiA* is reversibly induced and DdiA abundance reversibly increases population-wide independently of LexA and DdiA. Thus, DdiA neither positively nor negatively autoregulates *ddiA* expression.

DdiA, directly or indirectly, activates *dnaE2* transcription in MMC-treated and untreated cells. In MMC-treated cells, DdiA, directly or indirectly, also slightly but significantly inhibits *recX* transcription causing a decrease in RecX abundance. In untreated cells, *recX* transcript levels were similar in Δ*ddiA* and WT cells (Fig. 4F), while RecX was more abundant in the Δ*ddiA* cells (Fig. 4A). We suggest that DdiA also inhibits *recX* transcription in untreated cells but that this is not evident in the RT-qPCR analyses because *ddiA* expression is only activated in a minority of cells. Based on the increased RecQ abundance in untreated cells and those treated with MMC for 5hrs, we speculate, but have not shown, that DdiA also inhibits *recQ* transcription and RecQ abundance.

DnaE2 is an error-prone DNA polymerase that functions with the accessory factors ImuA and ImuB (57–60). Replicative DNA polymerases are highly accurate and processive, but they stall at most forms of DNA damage (4, 7). Since replication fork arrest is eventually lethal, cells need ways to cope with stalled DNA polymerases (4, 7). Because error-prone polymerases can incorporate any nucleotide opposite a replication-blocking DNA lesion and lack proofreading activity, they can carry out error-prone TLS across damaged DNA (4, 7). Once the damaged DNA has been successfully passed, the replicative DNA polymerase continues replication. While DdiA, directly or indirectly, activates *dnaE2* transcription, leading to an increase in DnaE2 abundance, ImuB was detected under all conditions, and its abundance increased by MMC treatment independently of DdiA. By contrast, ImuA was not detected under any condition tested. Because *imuA* and *imuB* are transcribed in an operon (47), we suggest that ImuA also accumulates independently of DdiA under all conditions and is not detected for technical reasons. The Δ*ddiA* mutant has a lower mutation frequency than WT in the absence of exogenous genotoxic stress. DnaE2, ImuA, and ImuB have been shown to contribute to mutagenesis in the absence of exogenous genotoxic stress in *C. crescentus* and *M. xanthus* (53, 54, 58) as well as to DNA damage-induced mutagenesis in *C. crescentus, M. tuberculosis,* and *M. xanthus* (53, 54, 57–59). Therefore, we suggest that the lower mutation frequency in the Δ*ddiA* mutant compared to WT in the absence of exogenous genotoxic stress is caused by the reduced *dnaE2* expression in the DdiA^+^ subpopulation. In response to exogenous genotoxic stress, DdiA and, therefore, DnaE2 abundance is increased population-wide, thereby enabling DnaE2/ImuA/ImuB TLS activity population-wide. Because DnaE2, ImuA, and ImuB contribute to DNA damage-induced mutagenesis in *M. xanthus* (53, 54), we predict but have not shown directly, that the Δ*ddiA* mutant also has a reduced mutation frequency in the presence of exogenous genotoxic stress because DnaE2 abundance is decreased. It has been argued that the activity of error-prone DNA polymerases represents a trade-off between fitness and mutagenesis (61). DdiA provides a fitness advantage in the absence of exogenous genotoxic stress. Because the lack of DdiA causes a lower mutation frequency, we suggest that DdiA mediates a trade-off between fitness and mutagenesis in the absence of exogenous genotoxic stress, and likely also in the presence of exogenous genotoxic stress. We propose that this trade-off is driven by the DdiA-dependent activation of *dnaE2* expression.

ImuB abundance (and by implication ImuA abundance) is equally upregulated in MMC-treated WT and Δ*ddiA* cells; however, the ImuB level is significantly lower in Δ*lexA* cells. Similarly, the abundance of the error-prone DNA polymerase DinB (also referred to as DNA polymerase IV) is increased by MMC stress independently of both LexA and DdiA. Altogether, the regulation of the abundance of ImuB and DinB implies that additional LexA- and DdiA-independent regulator(s), yet to be identified, is involved in regulating the accumulation of proteins involved in TLS.

RecX is a negative regulator of RecA that inhibits RecA recombinase activity and coprotease activity in *E. coli* (2, 41). In *E. coli*, *recX* is co-expressed with *recA* in a LexA-dependent manner, and it has been suggested that the increased RecX level contributes to turning off RecA activity and the LexA-dependent DDR (2, 41). In *M. xanthus*, RecX abundance is upregulated during MMC stress independently of LexA, and DdiA, directly or indirectly, inhibits *recX* transcription and, thus, RecX accumulation. These observations suggest that LexA- and DdiA-independent regulator(s) yet to be identified is involved in the upregulation of the RecX level in response to MMC treatment.

What, then, would be the logic of the LexA-independent and DdiA-dependent regulation of DnaE2 and RecX abundance? Because DnaE2 engages in error-prone TLS, we speculate that in the absence of exogenous genotoxic stress, a signal related to spontaneous replication stress caused by endogenous factors induces *ddiA* expression in a subpopulation of cells, but this signal is not sufficient to induce the RecA/LexA-dependent DDR. In this model, in the absence of exogenous genotoxic stress, the DdiA-dependent upregulation of DnaE2 would help to alleviate the replication stress by TLS. In parallel, the DdiA-dependent inhibition of RecX synthesis would increase the sensitivity of one or both RecA proteins to ssDNA. We speculate that the latter would be relevant in case the DnaE2-dependent TLS is insufficient to resolve the replication stress. In this model, the DdiA-dependent response is tailored to resolve replication stress. We speculate that an advantage of this tailored response to spontaneous replication stress could be that it is less costly than the induction of the complete RecA/LexA-dependent DDR in response to replication stress. Similarly, in response to exogenous genotoxic stress, the DdiA-dependent response would contribute to resolving replication stress. We speculate that the DdiA-dependent response contributes to generating the genetic variation that would help guarantee the survival of the *M. xanthus* population in the fluctuating terrestrial habitat.

LexA and DdrO are transcriptional repressors, proteolytically inactivated in response to DNA damage, and *lexA* and *ddrO* expression increases during the DDR due to negative autoregulation in the case of *lexA* and by an unknown mechanism in the case of *ddrO* (2, 3, 15, 26, 30). The binding of ssDNA by RecA activates the LexA co-protease activity, and similarly, the binding of ssDNA activates the PprI protease that cleaves DdrO (2, 3, 15, 30). The WYL domain-containing transcriptional activators PafBC, SiwR, and DriD are activated post-translationally in response to DNA damage by binding of ssDNA, and their abundance remains unchanged during the DDR (12, 18, 20, 23–25). The Ada-type transcriptional activators are activated post-translationally by DNA methylation damage, and their abundance increases upon activation due to positive autoregulation (1, 14, 33). Interestingly, the regulation of DdiA follows a different regulatory design, i.e., *ddiA* transcription and DdiA abundance are induced in response to DNA damage, but DdiA is not an autoregulator. Also, we have no evidence suggesting that DdiA is proteolytically cleaved or activated post-translationally. How *ddiA* expression is induced in response to DNA damage remains to be determined. In the future, it will be important to identify the signal and the mechanism for induction of *ddiA* expression. Similarly, it will be important to determine whether whether DdiA directly activates *dnaE2* and directly represses *recX* expression.

Phenotypic heterogeneity within a population of genetically identical bacterial cells has been suggested to be part of bet-hedging and/or a division of labor strategies that optimize the survival of the population (62–64). Generally, the diversification of cells into distinct subpopulations and the phenotype adopted by a particular cell are thought to be the result of stochastic processes (62, 63). We suggest that the heterogeneous activation of *ddiA* expression in the absence of exogenous genotoxic stress is neither part of such strategies nor stochastic. Rather as suggested this activation would be the result of spontaneous replication stress, which would subsequently be resolved by DnaE2/ImuA/ImuB.

Cells of the MMC-treated WT, the Δ*lexA* mutant, the Δ*ddiA* mutant, and the MMC-treated Δ*ddiA* mutant were significantly longer than untreated WT cells suggesting that DNA damage induces cell cycle checkpoint(s) impeding cell division in *M. xanthus*. Interference with chromosome replication and/or segregation inhibits cell division in *M. xanthus* (34, 37, 65). Therefore, based on the hypothesis that spontaneous replication stress induces *ddiA* expression and DdiA accumulation, we speculate that the cell division defect in the Δ*ddiA* mutant in the absence of exogenous genotoxic stress is caused by the blocked replication. In the future, it will be important to clarify how DNA damage inhibits cell division in a LexA-dependent manner in *M. xanthus*.

## Supporting information

All Supplementary Information

## Acknowledgment

We thank Dr. Maria Perez-Burgos, Dr. Anke Treuner-Lange, and Dr. Dominik Schumacher for many helpful discussions. This work was supported by the Max Planck Society.

## Conflict of Interest

The authors declare no conflict of interest.

## Availability of data and materials

The authors declare that all data supporting this study are available within the article and its Supplementary Information files. All materials used in the study are available from the corresponding author.

## Material & Methods

### Strains and cell growth

All *M. xanthus* strains used in this study are derivatives of the WT strain DK1622 (66) and are listed in Table S2. Plasmids and oligonucleotides are listed in Table S3 and Table S4, respectively. In-frame deletions were constructed by two-step homologous recombination as described (67). Plasmids were integrated in a single copy by site-specific recombination at the Mx8 *attB* site. All plasmids were verified by DNA sequencing, and all strains were verified by PCR. *M. xanthus* cultures were grown at 32°C in 1% CTT broth (1% [wt/vol] Bacto casitone, 10 mM Tris-HCl pH 8.0, 1 mM K_2_HPO_4_/KH_2_PO_4_ pH 7.6, 8 mM MgSO_4_) or on 1.5% agar supplemented with 1% CTT and kanamycin (50 μg mL^−1^) or oxytetracycline (10 μg mL^−1^) when appropriate (68). Growth in suspension culture was followed by measuring the optical density at 550nm (OD_550_). MMC and A22 were dissolved in 99.9% dimethyl sulfoxide (DMSO); ciprofloxacin and phleomycin in H_2_O, and nalidixic acid in 99.9% ethanol. Plasmids were propagated in *E. coli* NEB Turbo (F’ *proA^+^B^+^ lacI^q^ ΔlacZM15 /fhuA2 Δ(lac-proAB) glnV galK16 galE15 R(zgb-210::Tn10)*Tet^S^ *endA1 thi-1 Δ(hsdS-mcrB) (*New England Biolabs) at 37°C in lysogeny broth (LB) (69) supplemented with kanamycin (50 μg mL^−1^) or tetracycline (20 μg mL^−1^) when required.

### Cell length determination

5-µl aliquots of exponentially growing suspension cultures were spotted on 1% agarose supplemented with 0.2% CTT. Cells were immediately covered with a coverslip and imaged using a DMi8 inverted microscope and DFC9000 GT camera. To assess cell length, cells were segmented using Omnipose (70), segmentation was manually curated using Oufti (71), analyzed using Matlab R2020a (The MathWorks), and plotted using GraphPad Prism (GraphPad Software, LLC).

### Fluorescence microscopy

Fluorescence microscopy was performed as described (72). Briefly, exponentially growing cells were transferred to a 1.0% agarose pad (Cambrex) buffered with TPM buffer (10 mM Tris-HCl pH 7.6, 1 mM KPO_4_ pH 7.6, 8 mM MgSO_4_) and supplemented with 0.2% CTT broth on a microscope slide, and covered with a coverslip. A Leica DMi8 inverted microscope was used for imaging, and phase contrast and fluorescence images were acquired using a Hamamatsu ORCA-flash V2 Digital CMOS camera. For DAPI staining, cells were stained with 1mg mL^-1^ DAPI for 5min at 32°C. For image processing, Metamorph v 7.5 (Molecular Devices) was used.

### Immunoblot analysis

Immunoblots were performed as described (73). Rabbit polyclonal α-PilC (dilution: 1:2,000) (74) and α-mCh (dilution: 1:2,500) (BioVision) were used together with horseradish peroxidase-conjugated goat anti-rabbit immunoglobulin G (dilution: 1:10,000) (Sigma) as secondary antibody. Blots were developed using Luminata Forte Western HRP Substrate (Millipore) and visualized and quantified using a LAS-4000 luminescent image analyzer (Fujifilm). Proteins were separated by SDS-PAGE as described (73).

### RT-qPCR

Total RNA was isolated from exponentially growing *M. xanthus* strains in biological triplicates. Total RNA was extracted using the Monarch® Total RNA Miniprep Kit (New England Biolabs). Briefly, 10^9^ cells were harvested, resuspended in 200 μL lysis buffer (100 mM Tris-HCl pH 7.6, 1 mg mL^−1^ lysozyme), and incubated at 25°C for 5 min. The manufacturer’s protocol was followed to purify RNA. Next, Turbo DNase (Thermo Fisher Scientific) was added to the RNA following the manufacturer’s protocol and subsequently removed using the Monarch® RNA Cleanup Kit (50 μg; New England Biolabs). The LunaScript RT supermix kit (New England Biolabs) was used to generate complementary DNA (cDNA) using 1 μg RNA. qPCR was performed on the three biological replicates with each two technical replicates on an Applied Biosystems 7500 real-time PCR system using the Luna universal qPCR master mix (New England Biolabs) with the primers listed in Table S4. Differential gene expression analysis was performed following the comparative threshold cycle (*C_T_*) method (75). *MXAN_6066*, encoding TrpA, was used as an internal reference gene, as described (76). The *trpA* gene was used as a reference because the TrpA level was affected by neither MMC treatment nor lack of LexA or DdiA.

### Determination of the mutation frequency

Exponentially growing cultures in quadruplicates in 1% CTT broth were plated on 1.5% agar containing 1% CTT broth without (for viable cell counts) or with 25 μg mL^-1^ rifampicin to score Rif^R^ colony forming units. Mutation frequencies were calculated by dividing the number of Rif^R^ mutants by the number of cells analyzed in the experiments.

### Growth competition experiment

Exponentially growing cells of the WT and the Δ*ddiA* mutant in suspension culture were mixed at a 1:1 ratio and growth of the mixed culture followed by measuring OD_550_. Cells were kept in the exponential growth phase by repeated dilution. The ratio of the WT to the Δ*ddiA* mutant was measured using a qPCR approach with a primer pair (JJ53/JJ54) (Table S4) that amplified a DNA fragment across the *ddiA* gene, giving rise to DNA fragments with a length of 1749bp in the WT and 1345 bp in the Δ*ddiA* mutant. Specifically, chromosomal DNA was isolated from the co-culture immediately after mixing and then every 24hrs for a total of 120hrs. Subsequently, qPCR was performed with 15 cycles for amplification using Taq polymerase (VWR Life Science) and 0.2µg chromosomal DNA. In control experiments, we found that with 15 PCR cycles, the amplification remained in the exponential phase. The amplified DNA fragments were separated by agarose gel electrophoresis as described (73), visualized and quantified using a LAS-4000 luminescent image analyzer (Fujifilm), and the ratio between fragment abundance in the Δ*ddiA* mutant and WT calculated.

### Proteomic analysis using data independent acquisition-mass spectrometry (DIA-MS)

Whole-cell proteomics experiments were done using exponentially growing cultures in 1% CTT broth at 32°C of the indicated strains in suspension culture as described (77). For all strains analyzed, four biological replicates were analyzed. 35 mg of cells per sample were harvested and washed twice in 0.5 mL 1× phosphate-buffered saline (PBS) (137 mM NaCl, 2.7 mM KCl, 10 mM Na_2_HPO_4_, 1.8 mM KH_2_PO_4_, pH 7.5) supplemented with 2× protease inhibitor (Roche). The cells were sedimented and resuspended in 0.2 mL 0.1M ammonium bicarbonate containing 2% (w/vol) sodium lauroyl sarcosinate (SLS), followed by heat lysis at 95°C for 1hr. Next, the samples were centrifuged at 14,000× *g* for 5 min, and the supernatant was harvested. Next, 1.2 mL freezer-cold acetone was added to the supernatant, mixed, and incubated at −80°C for at least 2 hrs. Next, the samples were centrifuged at 21,000× *g* for 15 min at 4°C. The supernatant was discarded, and the pellet was washed three times with freezer-cold methanol. Next, the pellet was dried, and the methanol was completely removed. The protein pellet was resuspended in 200 μL 0.5% SLS (w/vol), and the protein amount was determined by bicinchoninic acid-based protein assay (Thermo Fisher Scientific). Proteins were reduced with 5 mM Tris(2-carboxyethyl) phosphine (Thermo Fisher Scientific) at 90°C for 15 min and alkylated using 10 mM iodoacetamide (Sigma Aldrich) at 25°C for 30 min in the dark. 50 µg protein was digested by 1 µg trypsin (Serva) at 30°C overnight. After digestion, SLS was precipitated by acidification, and peptides were desalted by using C18 solid phase extraction cartridges (Macherey-Nagel). Cartridges were prepared for sample loading by adding acetonitrile (ACN), followed by 0.1% trifluoroacetic acid (TFA) (Thermo Fisher Scientific). Peptides were loaded on equilibrated cartridges, washed with 5% ACN/0.1% TFA containing buffer, and finally eluted with 50% ACN and 0.1% TFA. Dried peptides were reconstituted in 0.1% TFA and then analyzed using Liquid Chromatography-Mass Spectrometry (LC-MS) using an Ultimate 3000 RSLC nano connected to an Exploris 480 Mass Spectrometer *via* a nanospray flex ion source (all Thermo FisherScientific) and an in-house packed HPLC C18 column (75μm × 42cm). The following separating gradient was used: 94% solvent A (0.15% formic acid) and 6% solvent B (99.85% ACN, 0.15% formic acid) to 25% solvent B over 65 minutes at a flow rate of 300 nL/min, followed by an additional increase of solvent B 35% over 24 min. MS raw data was acquired in data-independent acquisition mode. Briefly, spray voltage was set to 2.3 kV, the funnel radio frequency level at 40, and the ion transfer capillary heated to 275°C. For DIA experiments full MS resolutions were set to 120,000 at m/z 200 and full MS, AGC (Automatic Gain Control) target was 300% with an 50 ms IT (Ion Accumulation Time). Mass range was set to 350–1400. AGC target value for fragment spectra was set at 3000%. 45 windows of 14 Da plus 1 Da overlap were used. Resolution was set to 15,000 and MS/MS IT to 22 ms. Stepped HCD (high energy collision dissociation) collision energy of 25, 27.5, 30 % was used. MS1 data was acquired in profile, MS2 DIA data in centroid mode.

For analyzing DIA data, the neural network (NN) based DIA-NN suite version 1.8 (78) and an Uniprot protein database for *M. xanthus* were used. A data set centric spectral library for the DIA analysis was generated. DIA-NN performed noise interference correction (mass correction, RT prediction, and precursor/fragment co-elution correlation) and peptide precursor signal extraction of the DIA-NN raw data. The following parameters were used: Full tryptic digest was allowed with two missed cleavage sites, and oxidized methionines and carbamidomethylated cysteines as modifications. Match between runs and remove likely interferences were enabled. The NN classifier was set to the single-pass mode, and protein inference was based on genes. The quantification strategy was set to any LC (high accuracy). Cross-run normalization was set to RT-dependent. Library generation was set to smart profiling. DIA-NN outputs were further evaluated using the SafeQuant (79, 80) script modified to process DIA-NN outputs.

The mass spectrometry proteomics data of whole cell proteomics experiments have been deposited to the ProteomeXchange Consortium (81) *via* the PRIDE (82) partner repository with the dataset identifier PXD060688.

### Bioinformatics

Gene and protein sequences were obtained from the databases KEGG (83) and UniProt (84). The phylogenetic tree of myxobacteria was generated in MEGA-X (85) using the neighbor-joining method (86). Orthologs of an *M. xanthus* gene of interest in myxobacterial genomes were identified using the KEGG Sequence Similarity DataBase (83). The structure prediction of the DdiA dimer was performed with AlphaFold2-Multimer_v3 modeling *via* ColabFold (87, 88). To evaluate AlphaFold-generated models, predicted local distance difference test (pLDDT) and predicted alignment error (pAE) graphs of five models were made using a custom-made Matlab R2020a (The MathWorks) script. These models were ranked based on combined pLDDT and pAE values, with the best-ranked model used for further analysis and presentation. PyMOL (The PyMOL Molecular Graphics System, Version 2.4.1 Schrödinger, LLC) was used to analyze and visualize the structural model.

### Plasmid construction: pJJ34 (for generation of in-frame deletion of *ddiA*)

Up- and downstream fragments were amplified from genomic DNA of DK1622 using the primer pairs JJ13/JJ14 and JJ15/JJ16. Subsequently, the AB and CD fragments were used as templates for overlapping PCR with the primer pair JJ13/JJ16 to generate the AD fragment. The AD fragment was digested with HindIII + XbaI, and cloned in pBJ114. pJJ37 (for expression of P*_ddiA_*-*ddiA-mCh* from *attB*): the *ddiA* fragment was amplified with the primer pair JJ17/JJ20, and the mCherry fragment was amplified with the primer pair JJ21/JJ22 from pAH53 (37). Next, overlapping PCR was performed using the previous PCR products and the primer pair JJ17/JJ22. The product was digested with EcoRI and HindIII, and cloned into pSWU30. pJJ38 (replacement of *ddiA* with *ddiA*-*mCh* in the native site): up- and downstream fragments were amplified using pJJ37 as DNA template and the primer pairs JJ23/JJ24 and JJ25/JJ16. To generate the full-length insert, an overlapping PCR using the two fragments as DNA templates and the primer pair JJ23/JJ16 was performed. The fragment was digested with XbaI and HindIII, and cloned into pBJ114. pJJ47 (for generation of in-frame deletion of *lexA*): up- and downstream fragments were amplified using the primer pairs JJ45/J46 and JJ47/JJ48. Subsequently, the AB and CD fragments were used as templates for overlapping PCR with the primer pair JJ45/JJ48 to generate the AD fragment. The AD fragment was digested with HindIII and XbaI, and cloned in pBJ114. pJJ50 (for expression of P*_ddiA_*-*mCh* from *attB*): the P*_ddiA_* fragment was amplified with the primer pair JJ17/JJ58, and the *mCh* fragment was amplified with the primer pair JJ57/JJ32 from pJJ37. Next, overlapping PCR was performed using the previous PCR products and the primer pair JJ17/JJ32. The product was digested with EcoRI and HindIII, and cloned into pSW105. pJJ51 (for expression of P*_lexA_*-*lexA* from the *attB*): P*_lexA_*-*lexA* was amplified with the primer pair JJ59/JJ60. The fragment was digested with EcoRI and HindIII, and cloned in pSWU30.

## Notes

### Competing Interest Statement

The authors have declared no competing interest.

