## Supplementary material for "DdiA, an XRE family transcriptional regulator, regulates a LexA-independent DNA damage response in *Myxococcus xanthus*": All Supplementary Information

**This file contains:**

- Supplementary Figures 1-4
- Supplementary Tables 2-4
- Supplementary References

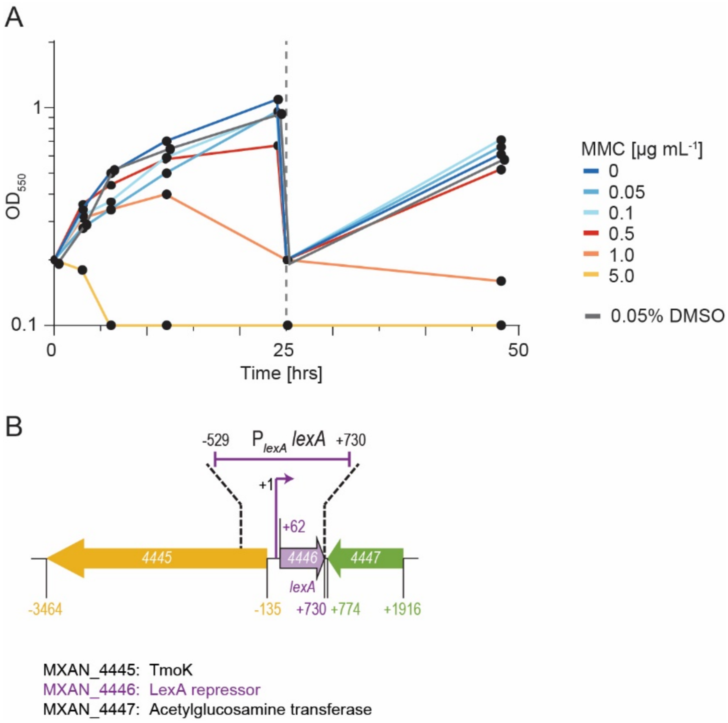


Figure S1. Determination of the highest non-lethal MMC concentration in *M. xanthus* and structure of *lexA* locus.
(A) Growth curves of WT *M. xanthus* in the presence of the indicated concentrations of MMC. Cells were grown in 1% CTT broth. Because MMC is dissolved in DMSO and the final DMSO concentration after addition to the growth medium is 0.05%, we also analyzed the growth of WT in the presence of 0.05% DMSO. At 24 hrs, the cultures were diluted into 1% CTT broth without MMC and DMSO. Data are from one representative experiment.
(B) Schematic of *lexA* locus. MXAN locus tags are included; genes are drawn to scale. +1 indicates the transcriptional start site of *lexA* (1). Numbers indicate the first and last nucleotide in start and stop codons, respectively relative to +1. Construct for the ectopic expression of *lexA* from its native promoter ($P_{lexA}$) is shown above. TmoK is a hybrid histidine protein kinase with a C-terminal GGDEF domain which lacks residues important for catalytic activity and c-di-GMP binding and is important for type IV pili-dependent motility (2).

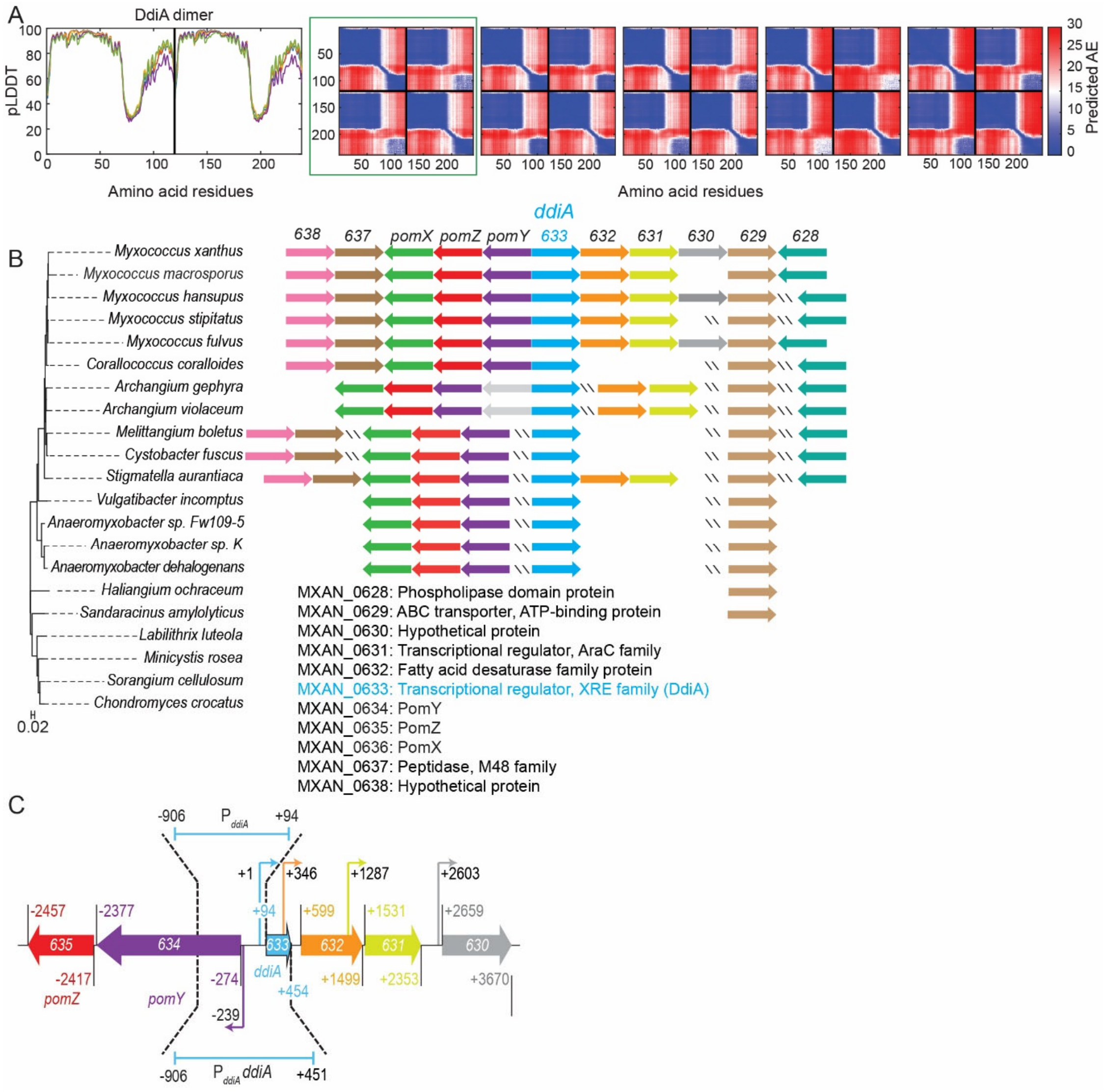


<u>Figure S2. Analysis of DdiA and the *ddiA* locus.</u>

(A) AlphaFold-Multimer structural model of DdiA dimer. Right panel, pLDDT plot for the five generated models of the DdiA dimer. Left panel, pAE plots for the five generated models of the DdiA dimer. The model highlighted by a green box was selected for further analysis.

(B) The *ddiA* locus is conserved in closely related myxobacteria. Transcription direction is indicated by the orientation of arrows with MXAN numbers indicated for the *ddiA* locus in *M. xanthus*.

(C) Detailed schematic of the *ddiA* locus. MXAN locus tags are included, genes are drawn to scale. +1 indicates the transcriptional start site of *ddiA* (1). Kinked arrows indicate transcriptional start sites relative to +1. Numbers indicate the first and last nucleotide in start and stop codons, respectively relative to +1. The blue bar labeled $P_{ddiA}$ indicates the fragment upstream of the *ddiA* start codon used to generate the $P_{ddiA}$-*mCh* promoter fusion. The blue bar labeled $P_{ddiA}$ *ddiA* indicates the fragment used for the ectopic expression of *ddiA-mCh* from the native $P_{ddiA}$.

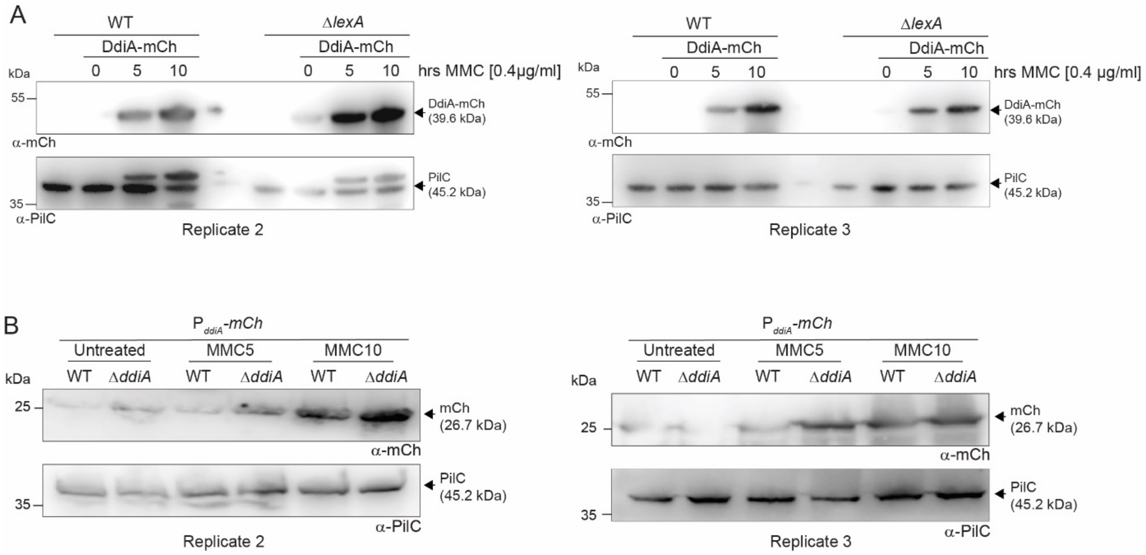


Figure S3. Analysis of DdiA-mCh abundance and $P_{ddiA}$ expression.

(A) Immunoblot analysis of DdiA-mCh abundance. Cells were grown and DdiA-mCh abundance was quantified as in Fig. 3E. The two blots represent the two replicates, which together with the replicate in Fig. 3E, were used to quantify DdiA-mCh by immunoblotting. (B) Immunoblot analysis of $P_{ddiA}$ expression. $P_{ddiA}$ was fused to a promoterless *mCh*. Cells were grown and mCh abundance was quantified as in Fig.3G. The two blots represent the two replicates, which together with the replicate in Fig.3G, were used to quantify mCh by immunoblotting.

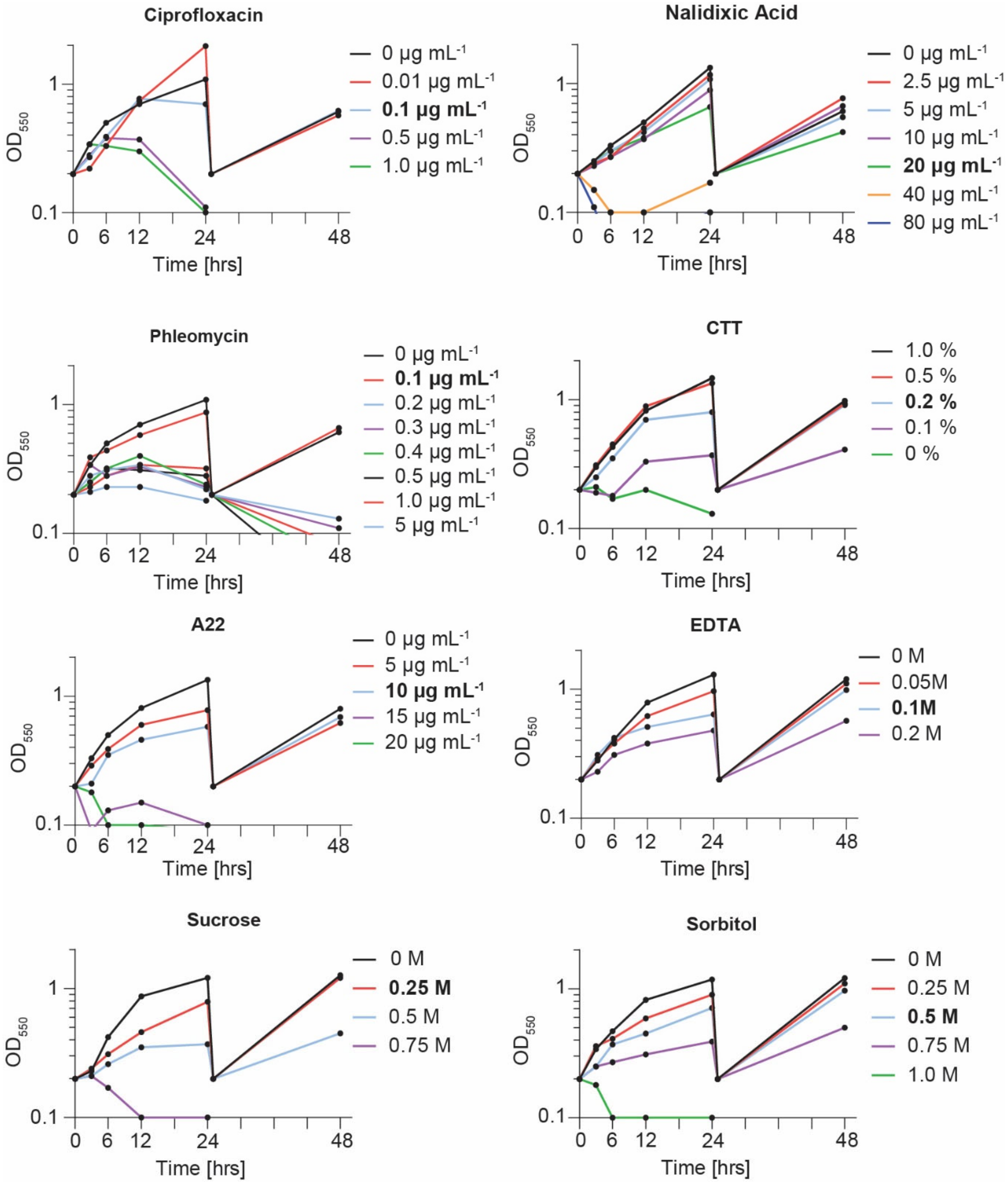


Figure S4. Growth curves of *M. xanthus* WT in the presence of various stressors. Growth curves of WT *M. xanthus* in the presence of the indicated stressors. Cells were grown in 1% CTT broth. At 24 hrs, the cultures were diluted into 1% CTT broth without the stressor. Data are from one representative experiment. Bold marks the conditions used in the experiments in Fig. 5D.

**Table S2.** *M. xanthus* strains used in this study.

| Strain | Genotype | Reference |
|---|---|---|
| DK1622 | Wildtype | (3) |
| SA9439 | Δ*lexA* | This study |
| SA9440 | Δ*lexA*, *attB*::pJJ51 ($P_{lexA}$ *lexA*) | This study |
| SA9405 | Δ*ddiA* | This study |
| SA9409 | Δ*ddiA*, *attB*::pJJ37 ($P_{ddiA}$ *ddiA-mCh*) | This study |
| SA9433 | *ddiA*::*ddiA-mCh* | This study |
| SA9438 | *ddiA*::*ddiA-mCh* Δ*lexA* | This study |
| SA9441 | *attB*::$P_{ddiA}$-*mCh* (pJJ50) | This study |
| SA9442 | *attB*::$P_{ddiA}$-*mCh* (pJJ50) Δ*ddiA* | This study |

**Table S3.** Plasmids used in this study.

| Plasmid | Description | Reference |
|---|---|---|
| pSW105 | *attP*, $P_{pilA}$, $Km^R$ | (4) |
| pSWU30 | *attP* $Tet^R$ | (5) |
| pBJ114 | *galK*; $Km^R$ | (6) |
| pJJ34 | pBJ114; for generation of in-frame deletion of *ddiA* (*MXAN_0633*) | This study |
| pJJ37 | For integration at *attB* and expression of *ddiA-mCh* from $P_{ddiA}$ | This study |
| pJJ38 | For integration of *ddiA-mCh* at the native site | This study |
| pJJ47 | pBJ114; For generation of in-frame deletion of *lexA* (*MXAN_4447*) | This study |
| pJJ50 | For integration at *attB* and expression of *mCh* from from $P_{ddiA}$ | This study |
| pJJ51 | For integration at *attB* and expression of *lexA* from $P_{lexA}$ | This study |

**Table S4.** Oligonucleotides used in this study.

| Name | Sequence ( 5´ to 3´) | Brief description |
|---|---|---|
| JJ13_DdiA_A_HindIII | GCGAAGCTTCTCCAGCTCCGCGACCTCACGGCGC | For construction of pJJ34 |
| JJ14_DdiA_B | CGCCACCTGCCCTCCAATGAGGGTTGCCAGTC | |
| JJ15_DdiA_C | CCCTCATTGGAGGGCAGGTGGCGAACGCGCT | |
| JJ16_DdiA_D_XbaI | GCGTCTAGATGCCAGACCTGGCCCAGCGTCTCG | |
| JJ53_DdiA_E | GGATGCGCCCCACGTCCAGCAG | |
| JJ54_DdiA_F | CAGCGCGGGCTTCCACACGATG | |
| JJ55_DdiA_G | CGCGGTCCGCGCGGCGCG | |
| JJ56_DdiA_H | GGCGAAGACGCGGTGCGTCC | |
| JJ45_LexA_A_HindIII | GCGAAGCTTGCTCCTTGTCCTTGCGCAGCTGCG | For construction of pJJ47 |
| JJ46_LexA_B | TGCGGCCGCCGCTCTTCCATGCGCTCCCTC | |
| JJ47_LexA_C | ATGGAAGAGCGGCGGCCGCACCCCGTAGTC | |
| JJ48_LexA_D_XbaI | GCGTCTAGACCACACCGCGGACGTGCTCG | |
| JJ49_LexA_E | GGACGAAGAGCGCCTCCACG | |
| JJ50_LexA_F | CCTCGAAGCAGCGCTACCCGG | |
| JJ51_LexA_G | GCGCGAGATTCTCAGCTTCATCG | |
| JJ52_LexA_H | CATCGACTGCCCCTTCACGC | |
| JJ17_EcoRI_up_633_fwd | GCGGAATTCCTGCAGGAAGCGCTGCGCCTCACGCGCC | For construction of pJJ37 |
| JJ20_633_Linker_rev | GGCGGCGGATCTGGCGGACGTCTCGAGCGCGTTCGCCACCTGC | |
| JJ21_Linker_mCh_fwd | TCCGCCAGATCCGCCGCCGGCTCCGGCATGGTGAGCAAGGGCG | |
| JJ22_HindIII_mCh_rev | GCGAAGCTTTTACTTGTACAGCTCGTCCATGCC | |
| JJ59_LexA_A_EcoRI | GCGGAATTCGCTCCTTGTCCTTGCGCAGCTGCG | For construction of pJJ51 |
| JJ60_LexA_rev_HindIII | CGCAAGCTTCTACGGGGTGCGGCCGCCCTGGAGC | |
| JJ23_HindIII_up_633_fwd | GCGAAGCTTCCGCGCAGGCCGAATTTTCGGCTTCCGCGC | For construction of pJJ38 |
| JJ24_down_0633_mCh_rev | CCGCTGTATCTTACTTGTACAGCTCGTCC | |
| JJ25_mCh_down_0633_fwd | GTACAAGTAAGATACAGCGGAGACCCGGCC | |
| JJ26_DdiA_D_XbaI | GCGTCTAGATGCCAGACCTGGCCCAGCGTCTCG | |
| JJ17_EcoRI_up_633_fwd | GCGGAATTCCTGCAGGAAGCGCTGCGCCTCACGCGCC | For construction of ppJJ50 |
| JJ57_PddiA_mCh_fwd | CCGCACGCCATGGTGAGCAAGGGCGAGGA | |
| JJ58_mCh_PddiA_rev | GCTCACCATGGCGTGCGGAAAACCTGTCG | |
| JJ59_HindIII_stop_mCh | GCGAAGCTTTTACTTGTACAGCTCGTCCATGCC | |
| JJ77_TrpA_fwd_3 | GATTGGCGTGGCGTTCAG | RT-qPCR for *trpA* |
| JJ78_TrpA_rev_3 | GGGGACATCGCTTGCGTA | |
| JJ81_UvrA_fwd_2 | ACAGATGGAGAAGCCGAAG | RT-qPCR for *uvrA* |
| JJ82_UvrA_rev_2 | ATAAAGCACCCGCAGGTAG | |
| JJ87_DdiA_fwd_2 | GGCAACCCTCATTGGAAACG | RT-qPCR for *ddiA* |
| JJ88_DdiA_rev_2 | CGTCCGACGAGCAGTTGA | |
| JJ91_DnaE2_fwd_1 | CGCCTCATTGGATGACTTG | RT-qPCR for *dnaE2* |
| JJ92_DnaE2_rev_1 | GCTCAATCTGCTTCACCAC | |
| JJ97_RecX_fwd_1 | AGGACCCGACAAGAAGAAGGAG | RT-qPCR for *recX* |
| JJ98_RecX_rev_1 | AACTTGATGGAGCGGTCCAC | |
